# Microstructural white matter and links with subcortical structures in chronic schizophrenia: A free-water imaging approach

**DOI:** 10.1101/621482

**Authors:** Tiril P. Gurholt, Unn K. Haukvik, Vera Lonning, Erik G. Jönsson, Ofer Pasternak, Ingrid Agartz

## Abstract

Schizophrenia is a severe mental disorder with often a chronic course. Neuroimaging studies report brain abnormalities in both white and gray matter structures. However, the relationship between microstructural white matter differences and volumetric subcortical structures is not known.

We investigated 30 long-term treated patients with schizophrenia (mean age 51.1±7.9 years, illness duration 27.6±8.0 years) and 42 healthy controls (mean age 54.1±9.1 years) using 3 T diffusion and structural magnetic resonance imaging. The free-water imaging method was used to model the diffusion signal, and subcortical volumes were obtained from FreeSurfer. We applied multiple linear regression to investigate associations between (i) patient status and regional white matter microstructure, (ii) medication dose or clinical symptoms on white matter microstructure in patients, and (iii) for interactions between subcortical volumes and diagnosis for microstructural white matter regions showing significant patient-control differences.

The patients had significantly decreased free-water corrected fractional anisotropy (FA_t_), explained by decreased axial diffusivity and increased radial diffusivity (RD_t_) bilaterally in the anterior corona radiata (ACR) and the left anterior limb of the internal capsule (ALIC) compared to controls. In the fornix, the patients had significantly increased RD_t_. In patients, positive symptoms were associated with localized increased free-water and negative symptoms with localized decreased FA_t_ and increased RD_t_. There were significant interactions between patient status and several subcortical structures on white matter microstructure and the free-water compartment for left ACR and fornix, and limited to the free-water compartment for right ACR and left ALIC. The Cohen’s d effect sizes were medium to large (0.61 to 1.20, absolute values).

The results suggest a specific pattern of frontal white matter axonal degeneration and demyelination and fornix demyelination that is attenuated in the presence of larger structures of the limbic system in patients with chronic schizophrenia. Findings warrants replication in larger samples.

## Introduction

Schizophrenia is a severe and often debilitating mental disorder with largely unknown disease mechanisms. It is well established that patients with schizophrenia, across different disease states, demonstrate white matter microstructural (1) and gray matter structural (2, 3) differences when compared to healthy controls, as well as progressive differences (4–6) related to the pathophysiology of the disorder and possibly medication use.

Diffusion magnetic resonance imaging (dMRI) and T1-weighted structural imaging are two popular magnetic resonance imaging (MRI) techniques that are often used to study schizophrenia. dMRI, using its popular analysis method - diffusion tensor imaging (DTI) (7) - yields in vivo indirect measures of white matter microstructure (8) such as fractional anisotropy (FA), axial diffusivity (AD) and radial diffusivity (RD). The FA measure is purported to be associated with white matter integrity, and can decrease both due to axonal degeneration and demyelination (6, 9), indicated by reduced AD and increased RD, respectively (8). However, the FA measure may not provide a good representation of white matter integrity due to several methodological issues (10), including partial volume effects e.g. from extracellular water contamination and crossing fibers (8). The bi-tensor free-water imaging model (11) accounts for extracellular free-water, yielding improved tissue specificity of white matter measures compared to the DTI model (11). The method also provides a free-water fractional volume measure that is affected by extracellular processes e.g. neuroinflammation, atrophy and cellular membrane breakdown (12).

The largest DTI meta-analysis to date showed that patients with schizophrenia have widespread white matter FA reductions compared to controls, with regionally more severe differences with increasing illness duration (1). Cross-sectional free-water imaging studies in schizophrenia corroborate increasing tissue change with illness duration; At schizophrenia onset, reports indicate limited tissue change together with widespread increase in free-water (13, 14), while with chronicity there is evidence of widespread tissue changes together with limited free-water increase (12, 15) when compared to healthy controls. These findings could indicate a severity gradient and that the temporal disease state needs to be considered in schizophrenia studies of microstructural white matter.

Structural MRI studies of patients with schizophrenia have shown widespread cortical thinning (3,16,17), area reduction (3, 17), folding abnormalities (18) and alterations of subcortical volumes, including smaller hippocampus and amygdala, and larger basal ganglia volumes (2,16,19), compared to healthy controls. Longitudinal studies have indicated, although somewhat inconclusively, increasing cortical thinning with illness duration (4). A recent large cross-sectional study showed enlargement of the putamen and pallidum volumes with age and illness duration (2). Microstructural white matter differences in schizophrenia indicated by reduced FA have been linked to cortical thinning in temporal regions, cuneus, frontal gyrus, orbitofrontal cortex (20), and cingulate cortex (21). Recently cortical thinning was also inversely associated with infracortical white matter anisotropy in adult patients (<50 years of age) (22). A single study showed increased mean diffusivity of the left accumbens, and the hippocampus and thalamus bilaterally, in patients (23). This could indicate patterns of associations between brain regions that are limited to patients with schizophrenia.

In the present study we investigated white matter diffusion properties using the free-water imaging method. The aims of the study were to: (i) identify differences in microstructural white matter diffusion properties between long-term treated patients with schizophrenia and healthy controls, (ii) investigate putative associations between medication or clinical symptoms on white matter microstructure in patients, and (iii) investigate whether volumetric measures of subcortical structures were associated with observed patient-control differences in white matter microstructure. In line with prior free-water imaging studies, we expected to observe microstructural white matter differences in patients with chronic schizophrenia together with limited evidence of increased free-water (12, 15), and clinical symptoms to be linked with microstructural white matter in patients (15). We hypothesized that white matter microstructure could be associated with medication use in line with prior DTI studies (24, 25), and that patient microstructural white matter differences are linked with volumetric subcortical measures, similar to prior cortical findings (20, 21).

## Materials and Methods

### Study Population

The subject sample consisted of 30 patients (Schizophrenia [N=22], Schizoaffective disorder [N=8]) and 42 controls, recruited among participants from the HUBIN study at the Karolinska Hospital (18, 26), and investigated between 2011 and 2015. Exclusion criteria for all participants were age <18 or >70 years, IQ<70, or previous severe head injury. All participants received oral and written information about the study and signed a written informed consent. The study was approved by the Research Ethics Committee at Karolinska Institutet, Sweden, and was conducted in accordance with the Helsinki declaration.

### Clinical Assessment

Patients and controls were assessed by a psychiatrist (EGJ) using the Structured Clinical Interview for DSM-III-R axis I disorders (27). Diagnosis was based on DSM-IV (28). Symptoms were assessed according to the Scale for the Assessment of Negative Symptoms (SANS) (29) and the Scale for the Assessment of Positive Symptoms (SAPS) (30). Psychosocial functioning was assessed using the split version of the Global Assessment of Function (GAF-S and GAF-F) scale (31). Age at onset was defined as the age of first verified positive psychotic symptom experience and duration of illness was calculated in years from age at onset to age at MRI. Chlorpromazine equivalent antipsychotic dose (CPZ) was computed (32).

### Data acquisition

Patients and controls underwent MRI on the same 3 T General Electric Healthcare Discovery MR750 Sigma scanner (General Electric Company, Milwaukee, Wisconsin, USA) equipped with an 8-channel head coil at the Karolinska Institutet and Hospital. The diffusion MRI data was acquired with 10 b_0_ volumes and 60 diffusion weighted volumes with b=1000 s/mm^2^. The scanning parameters were: 128×128 acquisition matrix, repetition time (TR)=6.0s, echo time (TE)=82.9ms, field of view (FOV)=240mm, flip angle=90° and spatial resolution 0.94×0.94×2.9mm^3^. A sagittal T1-weighted BRAVO sequence was acquired with TR=7.9s, TE=3.06s, inversion time (TI)=450ms, flip angle=12°, FOV=240mm and voxel size=0.94×0.94×1.2mm^3^. There was no major scanner upgrade or change of instrument during the study period.

### MRI Processing

All dMRI’s were processed as follows: Brain masks of the first b_0_ volume were manually edited to remove non-brain tissue. The dMRI’s were corrected for eddy current induced distortions and subject head motion using FSLs EDDY (33). We enabled automatic detection and correction of motion induced signal dropout (34), previously shown to enhance signal-to-noise ratio (35). EDDY outputs rotated b-vectors used in subsequent processing and total per-volume-movement used to calculate the average motion. Following EDDY correction, a bi-tensor diffusion model was fitted using a nonlinear regularized fit to obtain a free-water corrected diffusion tensor representing the tissue compartment and the fractional volume of an isotropic free-water compartment (11). From the diffusion tensor, a tissue specific scalar measurement of fractional anisotropy (FA_t_) was derived using FSLs dtifit. FA_t_ depends on two independent measures, radial diffusivity (RD_t_) and axial diffusivity (AD_t_), and they were derived for an additional level of investigation. The scalar measurements of each subject were projected onto a standard FA skeleton using Tract-Based Spatial Statistics (TBSS) (36). To do so, the FA images were registered to the ENIGMA-DTI FA template (37) that aligns with the Johns Hopkins University (JHU) DTI atlas (38) following the ENIGMA-DTI processing protocols (http://enigma.ini.usc.edu/protocols/dti-protocols/). Forty-four regions of interests (ROIs; Table 1) were extracted. The standard DTI measures of AD, RD and mean diffusivity (MD) were also derived and projected onto the FA skeleton.

**Table 1:** Overview of the investigated white matter ROIs.

| Abbreviation | Full name |
| --- | --- |
| *ACR\** | Anterior corona radiata |
| *ALIC\** | Anterior limb of internal capsule |
| *Average* | Average of diffusion measure |
| *BCC* | Body of corpus callosum |
| *CC* | Corpus callosum |
| *CGC\** | Cingulum |
| *CGH\** | Cingulum (hippocampal portion) |
| *CR\** | Corona Radiata |
| *CST\** | Corticospinal tract |
| *EC\** | External capsule |
| *FX* | Fornix |
| *FXST\** | Fornix stria terminalis |
| *GCC* | Genu of corpus callosum |
| *IC\** | Internal capsule |
| *IFO\** | Inferior fronto occipital fasciculus |
| *PCR\** | Posterior corona radiate |
| *PLIC\** | Posterior limb of internal capsule |
| *PTR\** | Posterior thalamic radiation |
| *RLIC\** | Retrolenticular part of IC |
| *SCC* | Splenium of corpus callosum |
| *SCR\** | Superior corona radiate |
| *SFO\** | Superior fronto-occipital fasciculus |
| *SLF\** | Superior longitudinal fasciculus |
| *SS\** | Sagittal stratum |
| *UNC\** | Uncinate |

\* Bilateral structures.

All T1-weighted MRI scans were processed using FreeSurfer (39) version 6.0.0. The processing steps include motion correction, bias field correction, brain extraction, intensity normalization and automatic Talairach transformation, with optimized 3T bias field filtration (40). Subcortical volumes were obtained through the subcortical segmentation stream (41), except for the hippocampus and amygdala structures obtained through joint segmentation in FreeSurfer version 6.0.0, development version (42, 43). The extracted bilateral subcortical structures were: hippocampus, amygdala, thalamus, nucleus accumbens, caudate, pallidum, putamen and lateral ventricle.

### MRI Quality control

Only dMRI’s and volumetric structures passing quality control were included in the analyses. Initially there were 80 participants in the study.

All DWIs were visually inspected in three orthogonal views for severe visible artifacts (44), leading to the exclusion of 4 participants. Additionally, we excluded 4 subjects with an EDDY estimated average motion above two standard deviations from the mean. After quality control there were 72 participants in the study.

All T1-weighted images were visually inspected for movement and cortical segmentation errors. No participants needed to be excluded at this stage. The segmentation quality of the subcortical volumes was assessed by manual inspection of outlier volumes (defined as: ≥1.5 times interquartile range). Outlier volumes were excluded if the segmentation was inaccurate. This led to the exclusion volumes from 4 participants, namely: one volume each for the left amygdala, left thalamus, left putamen, right putamen, right accumbens, and right caudate.

### Statistical method

The demographic variables of patients and controls were compared using χ^2^-test for categorical variables, two-sample t-test/two-sided Wilcoxon rank-sum test for normally/non-normally distributed continuous variables. Normality was assessed using the Shapiro-Wilk’s normality test (45).

In the main analyses, we applied multiple linear regression to assess the effect of patient-control differences on each white matter ROIs using the *lm*-function in R (version 3.5.0). For comparison, we included analyses using both the free-water imaging and standard DTI method. In patients, we further investigated free-water imaging metrics for the effects of medication and clinical symptoms on each ROI using similar models. For all models we adjusted for sex, age, and average movement.

We conducted follow-up analyses for ROIs showing significant patient-control differences to assess potential associations between white matter microstructure and volumetric measures of subcortical structures. To do so, we extended the main model to include a term for the subcortical volumes and its interaction with patient status. To capture associations between white matter microstructure and subcortical volumes, independent of size, we chose not to include ICV in the model.

We computed Cohen’s d effect size from the t-statistics for categorical variables, and via the partial correlation coefficient, r, for continuous variables (46). We corrected for multiple comparisons using the false discovery rate (FDR) at α=0.05 (47) across planed analyses, yielding significance threshold p≤0.0109. For follow-up analyses, a separate FDR threshold was computed at p≤0.0116.

## Results

### Demographic and Clinical Data

Patients had an average age of onset (AAO) of 23.5±4.6 years and an average duration of illness of 27.6±8.0 years. Of the patients, 93.3% received antipsychotic medication (8 first generation, 13 second generation, 7 first and second generation) with an average CPZ doze of 409.8±325.2 mg. Compared to the controls, patients had significantly fewer years of education (p=0.0011), and decreased functioning as assessed by GAF symptom (p=6.2e-13) and GAF function (p=1.4e-13) score. During diffusion MRI the patients moved significantly more than the controls (p=0.0298). There were no significant differences in the other clinical or demographic data (Table 2).

**Table 2:** Demographics and clinical variables.

| Clinical information<sup>a</sup> | Patients (N=30) | Healthy Controls (N=42) | $\chi^2$-test/Wilcoxon rank sum test/t-test | p-value |
| --- | --- | --- | --- | --- |
| Women, N (%) | 8 (26.7) | 13 (30.9) | 0.02 | 0.8954 |
| Age (years) | 51.1±7.9 | 54.1±9.1 | -1.43 | 0.1581 |
| Education (years) | 12.9±2.2 | 15.1±3.1 | -3.40 | **0.0011** |
| AAO (years) | 23.5±4.6 |  |  |  |
| DOI at MRI (years) | 27.6±8.0 |  |  |  |
| AP Medicated, N (%) | 28 (93.3) |  |  |  |
| FGA/SGA/mixed, N | 8/13/7 |  |  |  |
| CPZ (mg) | 409.8±325.2 |  |  |  |
| **Clinical measurements** |  |  |  |  |
| SAPS total | 9.2±7.9 |  |  |  |
| SANS total | 27.1±13.8 |  |  |  |
| GAF-S<sup>b</sup> | 46.5±10.0 | 81.7±9.0 | -35.00 | **6.2e-13** |
| GAF-F<sup>b</sup> | 45.5±8.9 | 86.7±7.6 | -40.00 | **1.4e-13** |

Notes: Significance threshold $p < 0.05$ indicated in bold. Two-sample t-test/two-sided Wilcoxon rank-sum test applied for normally/non-normally distributed continuous data. $\chi^2$ -test applied for categorical data. Abbreviations: AAO: Age at onset, AP: antipsychotic medication, CPZ: chlorpromazine equivalent antipsychotic dose, DOI: Duration of Illness, FGA: first generation antipsychotics, GAF: Global Assessment of Functioning, GAF-F: GAF function scale, GAF-S: GAF symptom scale, MRI: magnetic resonance imaging, Patients: schizophrenia patients, SAPS: Scale for the Assessment of Positive Symptoms, SANS: Scale for the Assessment of Negative Symptoms, SGA: second generation antipsychotics.
<sup>a</sup> Presented as mean±standard deviation, if not stated otherwise.
<sup>b</sup> Data not normally distributed.

### Patient-control differences in diffusion properties

FA_t_ was significantly lower in patients compared to controls in the right anterior corona radiata (ACR) (d=-0.96, p=0.0002), left ACR (d=-0.74, p=0.0040) and left anterior limb of internal capsule (ALIC) (d=-0.69, p=0.0069) (Figure 1; Table S1). In those regions, FA_t_ reductions were driven by significantly lower AD_t_ (d=-0.94, d=-0.75 and d=-0.71, respectively) and significantly higher RD_t_ (d=0.94, d=0.71, and d=0.70, respectively). Furthermore, the RD_t_ of the fornix was significantly higher for patients (d=0.82, p=0.0015) without any corresponding significant or nominal-significant differences in FA_t_ or AD_t_. There were nonsignificant patient-control differences in free-water.

**Figure 1:**
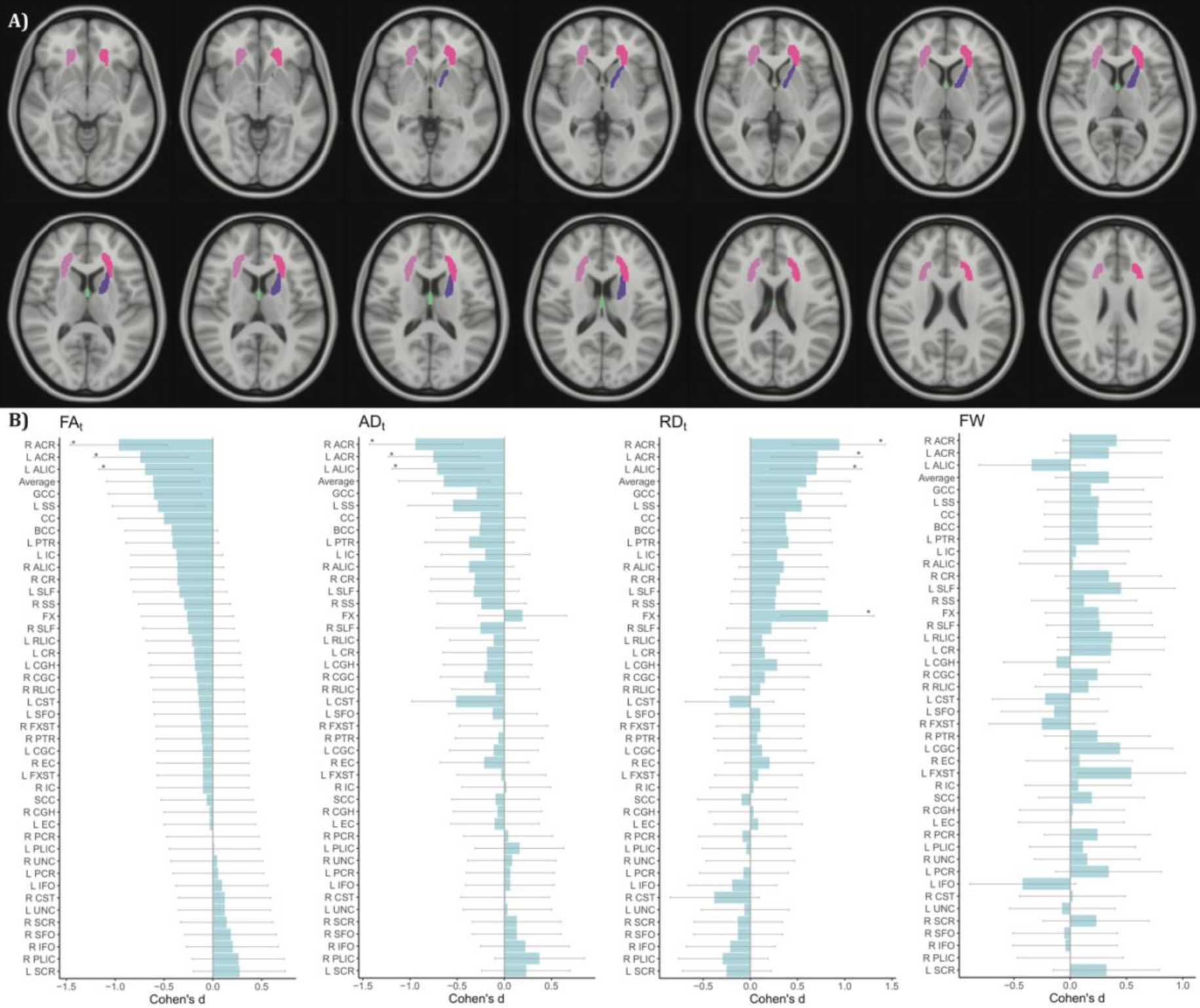
A) shows ROIs with significant patient-control differences are illustrated on the MNI 152 T1 atlas. B) shows Cohen’s d effect sizes for each ROI and free-water diffusion metric of patient-control differences. *Notes:* Ordered by ascending effect sizes for FA_t_. ROIs that pass the FDR threshold of p≤0.0109 are indicated with *. *Abbreviations:* ACR: anterior corona radiata, AD_t_: FW adjusted axial diffusivity, ALIC: anterior limb of internal capsule, Average: average of diffusion metric, BCC: body of corpus callosum, CC: corpus callosum, CGC: cingulum, CGH: cingulum hippocampal portion, CR: corona radiata, CST: corticospinal tract, EC: external capsule, FA_t_: FW adjusted fractional anisotropy, FW: free-water, FX: fornix, FXST: fornix stria terminalis, GCC: genu of corpus callosum, IC: internal capsule, IFO: inferior fronto occipital fasciculus, L: Left, PCR: posterior corona radiata, PLIC: posterior limb of internal capsule, PTR: posterior thalamic radiation, R: Right, RD_t_: FW adjusted Radial diffusivity, RLIC: retrolenticular part of IC, ROI: region of interest, SCC: splenium of corpus callosum, SCR: superior corona radiata, SFO: superior fronto-occipital fasciculus, SLF: superior longitudinal fasciculus, SS: sagittal stratum, UNC: uncinate.

For comparison, running the same analysis using the standard DTI model showed significant differences between patients and controls only for the right ACR (FA: d=-0.88; RD: d=0.66) (Table S2).

### Effects of medication

Analyses in patients did not show any significant CPZ medication effects on white matter microstructure (Table S3).

### Effects of clinical symptoms

Analyses in patients showed that total SAPS scores were significantly associated with increased free-water on the right posterior thalamic radiation (PTR; d=1.20, p=0.0061) and the left sagittal stratum (SS; d=1.16, p=0.0078), respectively (Table S4). Total SANS scores were for the right ALIC significantly associated with decreased FA_t_ (d=-1.14, p=0.0088) and increased RD_t_ (d=1.20, p=0.0061) (Table S5).

### Association between diffusion measures and subcortical volumes

In ROIs with significant diagnostic differences, we conducted follow-up analyses for possible association between free-water imaging diffusion metrics and subcortical volumes identified as significant interaction between subcortical volumes and patient status.

For the left ACR, the FA_t_ reduction in patients was attenuated in the presence of larger measures of hippocampus (left: d=0.77, p=0.0028; right: d=0.69, p=0.0068), right amygdala (d=0.67, p=0.0084), and right thalamus (d=0.68, p=0.0076). Similarly, AD_t_ reduction and RD_t_ increase were attenuated for larger left hippocampus (AD_t_: d=0.72, p=0.0048; RD_t_: d=-0.7, p=0.0064) and right thalamus (AD_t_: d=0.67, p=0.0084; RD_t_: d=-0.66, p=0.0098). For the free-water compartment, despite non-significant main effect of diagnosis, free-water was attenuated in the presence of larger caudate (left: d=-0.69, p=0.0069; right: d=-0.71, p=0.0057) and left thalamus (d=-0.69, p=0.0075) (Figure 2; Table S6).

**Figure 2:**
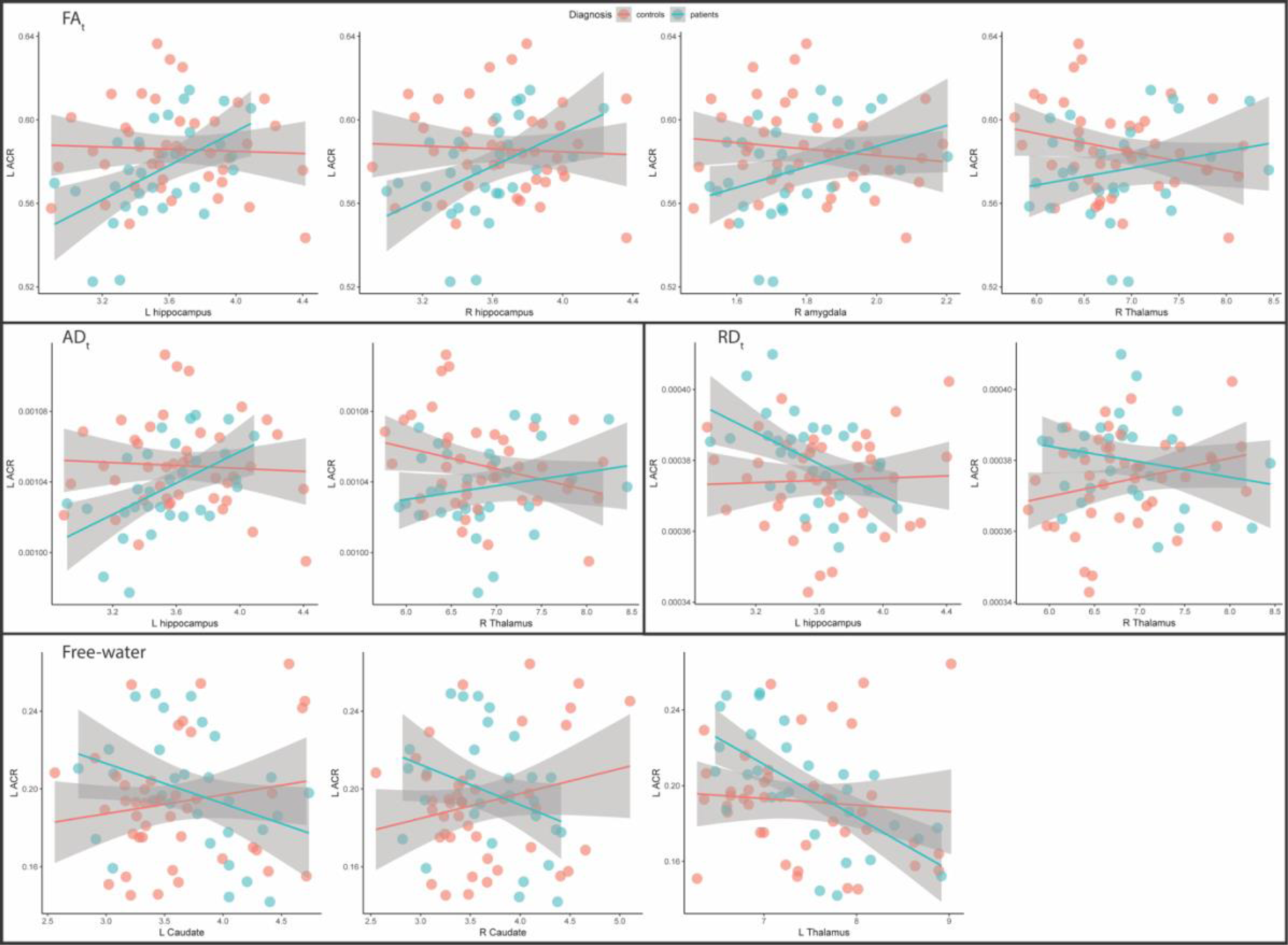
Scatterplots of the left anterior corona radiata with the subcortical structures that have significant interaction with diagnosis. *Notes:* Unadjusted locally weighted smoothing (LOESS) plots. *Abbreviations:* AD_t_: Free-water adjusted axial diffusivity, ACR: anterior corona radiata, FA_t_: Free-water adjusted fractional anisotropy, L: Left, RD_t_: Free-water adjusted Radial diffusivity, R: Right.

For the right ACR and left ALIC, despite non-significant main effect of diagnosis, we find associations between free-water and caudate in patients; the free-water compartment was reduced for the right ACR in the presence of larger caudate (left: d=-0.67, p=0.0091; right: d=-0.85, p=0.0012), and for the left ALIC for larger right caudate (d=-0.65, p=0.0111) (Figure 3; Table S7-S8).

**Figure 3:**
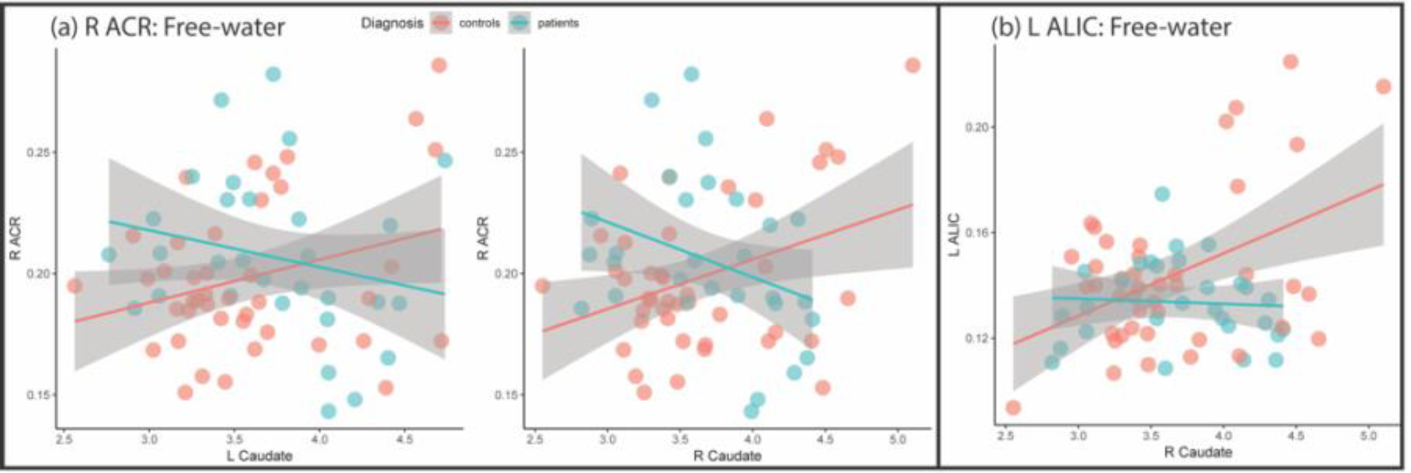
Scatterplots of the (a) right anterior corona radiata and (b) left anterior limb of internal capsule with the subcortical structures that have significant interaction with diagnosis. *Notes:* Unadjusted locally weighted smoothing (LOESS) plots. *Abbreviations:* ACR: anterior corona radiata, ALIC: anterior limb of internal capsule L: Left, R: Right.

For the fornix, RD_t_ was reduced for larger left nucleus accumbens (d=-0.73, p=0.0044) and hippocampus (left: d=-0.68, p=0.0083; right: d=-0.66, p=0.0100). The free-water compartment was, despite non-significant main effect of diagnosis, significantly associated with right ventricle (d=-0.69, p=0.0071) in patients, but this association appears driven by outliers (Figure 4; Table S9).

**Figure 4:**
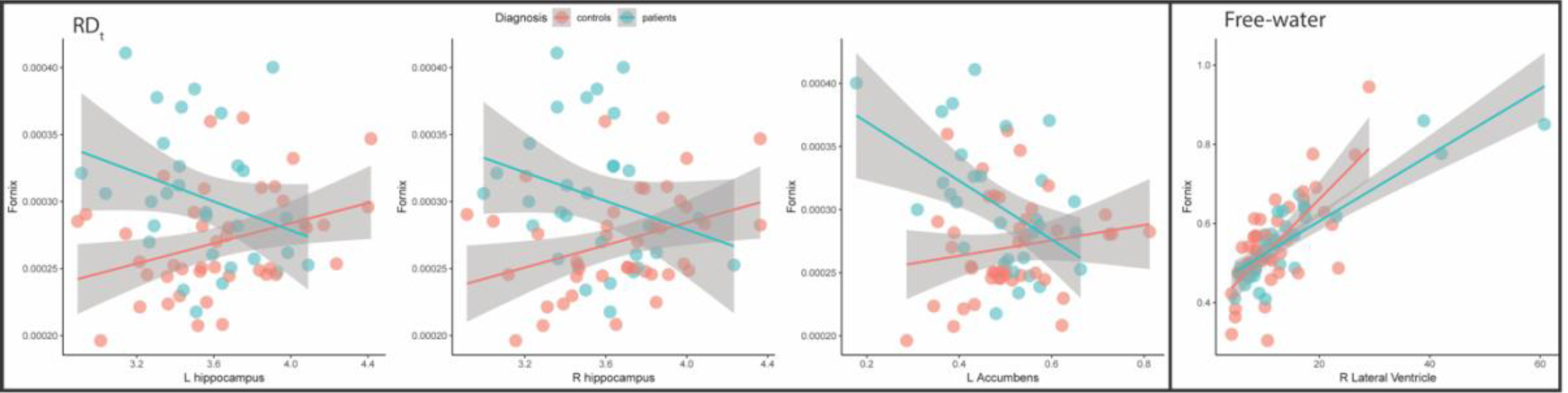
Scatterplots of the fornix with the subcortical structures that have significant interaction with diagnosis. *Notes:* Unadjusted locally weighted smoothing (LOESS) plots. *Abbreviations:* L: Left, RDt: Free-water adjusted Radial diffusivity, R: Right.

## Discussion

In this study we investigated the relationship between free-water imaging measures and subcortical volumes in patients with long-term treated chronic schizophrenia. The main findings were localized lower white matter anisotropy in patients compared with healthy controls, but no differences in free-water. White matter microstructure was linked with subcortical volumes in differing patterns for patients and controls.

The observed reduction in FA_t_, driven by a combination of AD_t_ reduction and RD_t_ increase, could indicate a pattern of axonal degeneration and demyelination in the frontal white matter in long-term treated chronic schizophrenia patients when compared to controls. In the fornix, the increased RD_t_ without simultaneous reduced FA_t_ and AD_t_ could imply demyelination without axonal degeneration in the patients. The free-water imaging method was more sensitive to patient-control differences in white matter microstructure than the standard DTI model. Our findings were similar to and partly overlapping with previous free-water imaging studies indicating reduced FA_t_ in cross-hemisphere frontal white matter in patients with schizophrenia (12–15), and no free-water increase in the chronic state (12, 15). The FA_t_ changes were more pronounced than the reported changes in first episode patients (13, 14). However, the FA_t_ changes were not as widespread as the previously reported changes in chronic schizophrenia (12, 15). The differences could be due to the limited sample size in the current study, but could also reflect that our long-term treated chronic schizophrenia sample on average had been ill for 27 years. This is longer than the previous studies, and it is not clearly known how the disorder progress with age and illness duration as captured by dMRI.

The observed interactions between patient status and subcortical structures on white matter microstructure for both the left ACR and the fornix, indicate patterns of association between the structures that are different in patients with chronic schizophrenia compared to controls. This is in line with previous studies that show cortical thinning in relation to white matter changes in patients with schizophrenia (20–22). The results may suggest that volumetric properties of brain anatomical structures are related to disrupted white matter microstructure in schizophrenia. The findings were in the direction of larger subcortical structures being associated with less severe white matter changes in patients with chronic schizophrenia, or vice versa. This could point towards a severity gradient in structural changes where less pronounced microstructural changes in patients have weaker diagnosis specific links to subcortical structures. They could also indicate disease specific patterns of associations between subcortical structures and microstructural white matter properties in chronic schizophrenia, and of disrupted functioning of the limbic system (48) and prefrontal connections (49). The reported findings are in line with the hypothesis of schizophrenia being a disorder of dysconnectivity (50–52). It is well established that the limbic system plays a role in schizophrenia, and subcortical structures of the limbic system have previously been reported as reduced in schizophrenia (2). The current study provides further support for limbic system involvement in the disorder together with links to white matter structures, which need to be further investigated.

We did not find any general patient-control differences in free-water. Despite this, in the follow-up analyses we found evidence of diagnosis specific involvement between free-water and some subcortical structures, and particularly larger caudate. Thus, there could be diagnosis specific association patterns between free-water and subcortical structures, but the implications of this is unknown. We can only speculate that the association could be linked to e.g. better functioning or reduced inflammatory state, but this needs to be further investigated. These findings warrants replication in larger samples.

Among patients, positive symptoms were significantly associated with increased free-water in the right PTR and left SS. Similarly, negative symptoms were associated with reduced FA_t_ and increased RD_t_ in the right ALIC. This is in line with a prior study indicating that positive symptoms are associated with increased free-water and negative symptoms with reduced FA_t_ (15). Furthermore, negative and positive symptoms were previously linked to changes in white matter microstructure using standard DTI (1). The association between microstructural white matter and clinical symptoms needs further investigation.

We did not observe any CPZ medication effects on white matter microstructure, which is in line with prior research (1). However, given the long-term treatment of the patients in the current study we cannot rule out that the observed effects on brain structure is due to antipsychotic medication, although not captured by CPZ. First and second generation antipsychotic medication could be differently involved with brain structure as previously shown for basal ganglia structures (53). How this relates to white matter microstructure needs to be addressed in a larger sample.

There are some limitations for the current study. The cross-sectional design makes it difficult to distinguish cause from effect. Moreover, although the effect sizes are strong, the limited sample size calls for replications in larger independent samples. Strengths of the current study includes a well characterized patient sample that has been characterized with research assessment by one psychiatrist for 12 years, 3 T high quality dMRI acquisition, validated and robust analysis methods.

*To conclude*, this study provides further evidence for white matter abnormalities, as well as evidence for altered involvement of subcortical structures with white matter microstructure, in patients with chronic schizophrenia when compared to healthy controls. The microstructural white matter differences indicate a process of frontal axonal degeneration and demyelination, and fornix demyelination in the patients. Both positive and negative symptoms could be influenced by free-water or microstructural tissue properties. The observed interaction between subcortical structures and patient status on white matter microstructure could indicate disease specific patterns of associations between the structures, limited to patients. To fully capture the linkage between gray and white matter tissue in chronic schizophrenia, future studies are needed.

## Acknowledgements

We thank the study participants and clinicians involved in recruitment and clinical assessment at Karolinska Institutet, Sweden, and Sara Holmqvist, Charlotta Leandersson, Erik Söderman, Rosland Sitnikov (Karolinska Institutet, Sweden), Maja Hjorth-Johansen (University of Oslo, Norway), Stener Nerland (Diakonhjemmet Hospital/University of Oslo, Norway) and Francesco Bettella (Oslo University Hospital, Norway), for their assistance.

## Author contributions

TPG designed the study in collaboration with OP and IA. TPG did the literature search and drafted the manuscript and interpreted the data together with OP, UKH and VL. OP contributed with the free-water imaging method. EGJ and IA obtained funding, contributed to data acquisition and study design. All authors contributed to and approved the final manuscript.

## References

1. Kelly S, Jahanshad N, Zalesky A, Kochunov P, Agartz I, Alloza C, Andreassen OA, Arango C, Banaj N, Bouix S, et al. Widespread white matter microstructural differences in schizophrenia across 4322 individuals: results from the ENIGMA Schizophrenia DTI Working Group. Mol Psychiatry (2017) doi:10.1038/mp.2017.170 https://www.nature.com/articles/mp2017170#supplementary-information

2. van Erp TG, Hibar DP, Rasmussen JM, Glahn DC, Pearlson GD, Andreassen OA, Agartz I, Westlye LT, Haukvik UK, Dale AM, et al. Subcortical brain volume abnormalities in 2028 individuals with schizophrenia and 2540 healthy controls via the ENIGMA consortium. Mol Psychiatry (2016) 21:585. doi:10.1038/mp.2015.118

3. van Erp TGM, Walton E, Hibar DP, Schmaal L, Jiang W, Glahn DC, Pearlson GD, Yao N, Fukunaga M, Hashimoto R, et al. Cortical Brain Abnormalities in 4474 Individuals With Schizophrenia and 5098 Control Subjects via the Enhancing Neuro Imaging Genetics Through Meta Analysis (ENIGMA) Consortium. Biol Psychiatry (2018) doi:10.1016/j.biopsych.2018.04.023

4. Dietsche B, Kircher T, Falkenberg I. Structural brain changes in schizophrenia at different stages of the illness: A selective review of longitudinal magnetic resonance imaging studies. Aust N Z J Psychiatry (2017) 51:500–508. doi:10.1177/0004867417699473

5. Kochunov P, Hong LE. Neurodevelopmental and Neurodegenerative Models of Schizophrenia: White Matter at the Center Stage. Schizophr Bull (2014) 40:721–728. doi:10.1093/schbul/sbu070

6. Samartzis L, Dima D, Fusar-Poli P, Kyriakopoulos M. White Matter Alterations in Early Stages of Schizophrenia: A Systematic Review of Diffusion Tensor Imaging Studies. J Neuroimaging (2014) 24:101–110. doi:10.1111/j.1552-6569.2012.00779.x

7. Basser PJ, Mattiello J, LeBihan D. MR diffusion tensor spectroscopy and imaging. Biophys J (1994) 66:259–67. doi:10.1016/S0006-3495(94)80775-1

8. Alexander AL, Lee JE, Lazar M, Field AS. Diffusion tensor imaging of the brain. Neurotherapeutics (2007) 4:316–29. doi:10.1016/j.nurt.2007.05.011

9. Mighdoll MI, Tao R, Kleinman JE, Hyde TM. Myelin, myelin-related disorders, and psychosis. Schizophr Res (2015) 161:85–93. doi:http://dx.doi.org/10.1016/j.schres.2014.09.040

10. Jones DK, Knösche TR, Turner R. White matter integrity, fiber count, and other fallacies: the do’s and don’ts of diffusion MRI. NeuroImage (2013) 73:239–254. doi:10.1016/j.neuroimage.2012.06.081

11. Pasternak O, Sochen N, Gur Y, Intrator N, Assaf Y. Free water elimination and mapping from diffusion MRI. Magn Reson Med (2009) 62:717–30. doi:10.1002/mrm.22055

12. Oestreich LKL, Lyall AE, Pasternak O, Kikinis Z, Newell DT, Savadjiev P, Bouix S, Shenton ME, Kubicki M, Australian Schizophrenia Research B, et al. Characterizing white matter changes in chronic schizophrenia: A free-water imaging multi-site study. Schizophr Res (2017) 189:153–161. doi:10.1016/j.schres.2017.02.006

13. Pasternak O, Westin CF, Bouix S, Seidman LJ, Goldstein JM, Woo TU, Petryshen TL, Mesholam-Gately RI, McCarley RW, Kikinis R, et al. Excessive extracellular volume reveals a neurodegenerative pattern in schizophrenia onset. J Neurosci (2012) 32:17365–72. doi:10.1523/JNEUROSCI.2904-12.2012

14. Lyall AE, Pasternak O, Robinson DG, Newell D, Trampush JW, Gallego JA, Fava M, Malhotra AK, Karlsgodt KH, Kubicki M, et al. Greater extracellular free-water in first-episode psychosis predicts better neurocognitive functioning. Mol Psychiatry (2018) 23:701–707. doi:10.1038/mp.2017.43

15. Pasternak O, Westin CF, Dahlben B, Bouix S, Kubicki M. The extent of diffusion MRI markers of neuroinflammation and white matter deterioration in chronic schizophrenia. Schizophr Res (2015) 161:113–8. doi:10.1016/j.schres.2014.07.031

16. Rimol LM, Hartberg CB, Nesvag R, Fennema-Notestine C, Hagler DJ, Pung CJ, Jennings RG, Haukvik UK, Lange E, Nakstad PH, et al. Cortical thickness and subcortical volumes in schizophrenia and bipolar disorder. Biol Psychiatry (2010) 68:41–50. doi:10.1016/j.biopsych.2010.03.036

17. Rimol LM, Nesvag R, Hagler DJ, Bergmann O, Fennema-Notestine C, Hartberg CB, Haukvik UK, Lange E, Pung CJ, Server A, et al. Cortical volume, surface area, and thickness in schizophrenia and bipolar disorder. Biol Psychiatry (2012) 71:552–60. doi:10.1016/j.biopsych.2011.11.026

18. Nesvåg R, Schaer M, Haukvik UK, Westlye LT, Rimol LM, Lange EH, Hartberg CB, Ottet M-C, Melle I, Andreassen OA, et al. Reduced brain cortical folding in schizophrenia revealed in two independent samples. Schizophr Res (2014) 152:333–338. doi:https://doi.org/10.1016/j.schres.2013.11.032

19. Haukvik UK, Westlye LT, Morch-Johnsen L, Jorgensen KN, Lange EH, Dale AM, Melle I, Andreassen OA, Agartz I. In vivo hippocampal subfield volumes in schizophrenia and bipolar disorder. Biol Psychiatry (2015) 77:581–8. doi:10.1016/j.biopsych.2014.06.020

20. Ehrlich S, Geisler D, Yendiki A, Panneck P, Roessner V, Calhoun VD, Magnotta VA, Gollub RL, White T. Associations of white matter integrity and cortical thickness in patients with schizophrenia and healthy controls. Schizophr Bull (2014) 40:665–74. doi:10.1093/schbul/sbt056

21. Koch K, Schultz CC, Wagner G, Schachtzabel C, Reichenbach JR, Sauer H, Schlosser RG. Disrupted white matter connectivity is associated with reduced cortical thickness in the cingulate cortex in schizophrenia. Cortex J Devoted Study Nerv Syst Behav (2013) 49:722–9. doi:10.1016/j.cortex.2012.02.001

22. Di Biase MA, Cropley VL, Cocchi L, Fornito A, Calamante F, Ganella EP, Pantelis C, Zalesky A. Linking Cortical and Connectional Pathology in Schizophrenia. Schizophr Bull (2018) doi:10.1093/schbul/sby121

23. Spoletini I, Cherubini A, Banfi G, Rubino IA, Peran P, Caltagirone C, Spalletta G. Hippocampi, Thalami, and Accumbens Microstructural Damage in Schizophrenia: A Volumetry, Diffusivity, and Neuropsychological Study. Schizophr Bull (2009) 37:118–130. doi:10.1093/schbul/sbp058

24. Ozcelik-Eroglu E, Ertugrul A, Oguz KK, Has AC, Karahan S, Yazici MK. Effect of clozapine on white matter integrity in patients with schizophrenia: a diffusion tensor imaging study. Psychiatry Res (2014) 223:226–235. doi:10.1016/j.pscychresns.2014.06.001

25. Ebdrup BH, Raghava JM, Nielsen MØ, Rostrup E, Glenthøj B. Frontal fasciculi and psychotic symptoms in antipsychotic-naive patients with schizophrenia before and after 6 weeks of selective dopamine D2/3 receptor blockade. J Psychiatry Neurosci JPN (2016) 41:133–141.

26. Ekholm B, Ekholm A, Adolfsson R, Vares M, Ösby U, Sedvall GC, Jönsson EG. Evaluation of diagnostic procedures in Swedish patients with schizophrenia and related psychoses. Nord J Psychiatry (2005) 59:457–464. doi:10.1080/08039480500360906

27. Spitzer RL, Williams JBW, Gibbon M, First MB. Structured Clinical Interview for DSM-III-R - Patient Version (SCID-P). (1988)

28. American Psychiatric Association. Diagnostic and Statistical Manual of Mental Disorders, International Version. Fourth. Washington DC: American Psychiatric Association (1995).

29. N. C. Andreasen. The Scale for the Assessment of Negative Symptoms (SANS). University of Iowa, Iowa City, IA (1983).

30. N. C. Andreasen. The Scale for the Assessment of Positive Symptoms (SAPS). University of Iowa, Iowa City, IA (1984).

31. Pedersen G, Hagtvet KA, Karterud S. Generalizability studies of the Global Assessment of Functioning-Split version. Compr Psychiatry (2007) 48:88–94. doi:10.1016/j.comppsych.2006.03.008

32. Woods SW. Chlorpromazine equivalent doses for the newer atypical antipsychotics. J Clin Psychiatry (2003) 64:663–7.

33. Andersson JLR, Sotiropoulos SN. An integrated approach to correction for off-resonance effects and subject movement in diffusion MR imaging. NeuroImage (2016) 125:1063–1078. doi:10.1016/j.neuroimage.2015.10.019

34. Andersson JLR, Graham MS, Zsoldos E, Sotiropoulos SN. Incorporating outlier detection and replacement into a non-parametric framework for movement and distortion correction of diffusion MR images. NeuroImage (2016) 141:556–572. doi:10.1016/j.neuroimage.2016.06.058

35. Tonnesen S, Kaufmann T, Doan NT, Alnaes D, Cordova-Palomera A, Meer DV, Rokicki J, Moberget T, Gurholt TP, Haukvik UK, et al. White matter aberrations and age-related trajectories in patients with schizophrenia and bipolar disorder revealed by diffusion tensor imaging. Sci Rep (2018) 8:14129. doi:10.1038/s41598-018-32355-9

36. Smith SM, Jenkinson M, Johansen-Berg H, Rueckert D, Nichols TE, Mackay CE, Watkins KE, Ciccarelli O, Cader MZ, Matthews PM, et al. Tract-based spatial statistics: voxelwise analysis of multi-subject diffusion data. NeuroImage (2006) 31:1487–505. doi:10.1016/j.neuroimage.2006.02.024

37. Jahanshad N, Kochunov PV, Sprooten E, Mandl RC, Nichols TE, Almasy L, Blangero J, Brouwer RM, Curran JE, de Zubicaray GI, et al. Multi-site genetic analysis of diffusion images and voxelwise heritability analysis: a pilot project of the ENIGMA-DTI working group. NeuroImage (2013) 81:455–69. doi:10.1016/j.neuroimage.2013.04.061

38. Mori S, Oishi K, Jiang H, Jiang L, Li X, Akhter K, Hua K, Faria AV, Mahmood A, Woods R, et al. Stereotaxic white matter atlas based on diffusion tensor imaging in an ICBM template. NeuroImage (2008) 40:570–582. doi:10.1016/j.neuroimage.2007.12.035

39. Fischl B. FreeSurfer. NeuroImage (2012) 62:774–81. doi:10.1016/j.neuroimage.2012.01.021

40. Zheng W, Chee MW, Zagorodnov V. Improvement of brain segmentation accuracy by optimizing non-uniformity correction using N3. NeuroImage (2009) 48:73–83. doi:10.1016/j.neuroimage.2009.06.039

41. Fischl B, Salat DH, Busa E, Albert M, Dieterich M, Haselgrove C, van der Kouwe A, Killiany R, Kennedy D, Klaveness S, et al. Whole brain segmentation: automated labeling of neuroanatomical structures in the human brain. Neuron (2002) 33:341–55. doi:http://dx.doi.org/10.1016/S0896-6273(02)00569-X

42. Iglesias JE, Augustinack JC, Nguyen K, Player CM, Player A, Wright M, Roy N, Frosch MP, McKee AC, Wald LL, et al. A computational atlas of the hippocampal formation using ex vivo, ultra-high resolution MRI: Application to adaptive segmentation of in vivo MRI. NeuroImage (2015) 115:117–37. doi:10.1016/j.neuroimage.2015.04.042

43. Saygin ZM, Kliemann D, Iglesias JE, van der Kouwe AJW, Boyd E, Reuter M, Stevens A, Van Leemput K, McKee A, Frosch MP, et al. High-resolution magnetic resonance imaging reveals nuclei of the human amygdala: manual segmentation to automatic atlas. NeuroImage (2017) 155:370–382. doi:10.1016/j.neuroimage.2017.04.046

44. Tournier JD, Mori S, Leemans A. Diffusion tensor imaging and beyond. Magn Reson Med (2011) 65:1532–56. doi:10.1002/mrm.22924

45. Royston P. Remark AS R94: A Remark on Algorithm AS 181: The W-test for Normality. Appl Stat (1995) 44:547–551. doi:10.2307/2986146

46. Nakagawa S, Cuthill IC. Effect size, confidence interval and statistical significance: a practical guide for biologists. Biol Rev Camb Philos Soc (2007) 82:591–605. doi:10.1111/j.1469-185X.2007.00027.x

47. Benjamini Y, Hochberg Y. Controlling the False Discovery Rate - a Practical and Powerful Approach to Multiple Testing. J R Stat Soc Ser B-Methodol (1995) 57:289–300.

48. Catani M, Dell’acqua F, Thiebaut de Schotten M. A revised limbic system model for memory, emotion and behaviour. Neurosci Biobehav Rev (2013) 37:1724–37. doi:10.1016/j.neubiorev.2013.07.001

49. Barbas H. General cortical and special prefrontal connections: principles from structure to function. Annu Rev Neurosci (2015) 38:269–289. doi:10.1146/annurev-neuro-071714-033936

50. Pettersson-Yeo W, Allen P, Benetti S, McGuire P, Mechelli A. Dysconnectivity in schizophrenia: Where are we now? Neurosci Biobehav Rev (2011) 35:1110–1124. doi:https://doi.org/10.1016/j.neubiorev.2010.11.004

51. van den Heuvel MP, Fornito A. Brain networks in schizophrenia. Neuropsychol Rev (2014) 24:32–48. doi:10.1007/s11065-014-9248-7

52. Schmitt A, Hasan A, Gruber O, Falkai P. Schizophrenia as a disorder of disconnectivity. Eur Arch Psychiatry Clin Neurosci (2011) 261:150. doi:10.1007/s00406-011-0242-2

53. Jørgensen KN, Nesvåg R, Gunleiksrud S, Raballo A, Jönsson EG, Agartz I. First- and second-generation antipsychotic drug treatment and subcortical brain morphology in schizophrenia. Eur Arch Psychiatry Clin Neurosci (2016) 266:451–460. doi:10.1007/s00406-015-0650-9

