## Supplementary information for "Microstructural white matter and links with subcortical structures in chronic schizophrenia: A free-water imaging approach"

*Table S1:* The Cohen's d effect size of patient-control differences for $FA_t$, $AD_t$, $RD_t$ and FW.

| | **$FA_t$** | | | **$AD_t$** | | | **$RD_t$** | | | **FW** | | |
|---|---|---|---|---|---|---|---|---|---|---|---|---|
| **ROI** | **d** | **t** | **p** | **d** | **t** | **p** | **d** | **t** | **p** | **d** | **t** | **p** |
| **R ACR** | −0.96 | −3.88 | **0.0002** | −0.94 | −3.78 | **0.0003** | 0.94 | 3.79 | **0.0003** | 0.41 | 1.65 | 0.1038 |
| **L ACR** | −0.74 | −2.98 | **0.0040** | −0.75 | −3.02 | **0.0035** | 0.71 | 2.87 | **0.0055** | 0.34 | 1.37 | 0.1738 |
| **L ALIC** | −0.69 | −2.79 | **0.0069** | −0.71 | −2.85 | **0.0058** | 0.70 | 2.81 | **0.0064** | −0.34 | −1.37 | 0.1748 |
| **Average** | −0.61 | −2.48 | 0.0156 | −0.64 | −2.57 | 0.0124 | 0.59 | 2.36 | 0.0211 | 0.34 | 1.39 | 0.1685 |
| **GCC** | −0.60 | −2.40 | 0.0189 | −0.29 | −1.16 | 0.2521 | 0.49 | 1.98 | 0.0514 | 0.18 | 0.71 | 0.4784 |
| **L SS** | −0.56 | −2.24 | 0.0281 | −0.54 | −2.17 | 0.0332 | 0.54 | 2.16 | 0.0343 | 0.25 | 1.03 | 0.3089 |
| **CC** | −0.50 | −2.00 | 0.0496 | −0.25 | −1.01 | 0.3147 | 0.37 | 1.49 | 0.1401 | 0.24 | 0.97 | 0.3342 |
| **BCC** | −0.42 | −1.70 | 0.0932 | −0.26 | −1.04 | 0.3020 | 0.38 | 1.53 | 0.1300 | 0.24 | 0.99 | 0.3268 |
| **L PTR** | −0.41 | −1.66 | 0.1009 | −0.37 | −1.49 | 0.1414 | 0.40 | 1.61 | 0.1118 | 0.25 | 1.02 | 0.3136 |
| **L IC** | −0.37 | −1.50 | 0.1392 | −0.20 | −0.82 | 0.4149 | 0.28 | 1.13 | 0.2622 | 0.05 | 0.22 | 0.8281 |
| **R ALIC** | −0.36 | −1.47 | 0.1458 | −0.37 | −1.49 | 0.1414 | 0.35 | 1.42 | 0.1610 | 0.02 | 0.07 | 0.9404 |
| **R CR** | −0.36 | −1.47 | 0.1469 | −0.31 | −1.24 | 0.2176 | 0.31 | 1.23 | 0.2222 | 0.34 | 1.36 | 0.1777 |
| **L SLF** | −0.34 | −1.35 | 0.1806 | −0.32 | −1.29 | 0.2001 | 0.27 | 1.11 | 0.2721 | 0.45 | 1.84 | 0.0709 |
| **R SS** | −0.29 | −1.19 | 0.2400 | −0.24 | −0.97 | 0.3333 | 0.26 | 1.03 | 0.3045 | 0.12 | 0.59 | 0.6164 |
| **FX** | −0.26 | −1.06 | 0.2933 | 0.19 | 0.77 | 0.4449 | 0.82 | 3.32 | **0.0015** | 0.25 | 1.03 | 0.3083 |
| **R SLF** | −0.25 | −1.02 | 0.3118 | −0.25 | −1.00 | 0.3220 | 0.22 | 0.88 | 0.3803 | 0.26 | 1.04 | 0.3022 |
| **L RLIC** | −0.21 | −0.84 | 0.4041 | −0.11 | −0.43 | 0.6693 | 0.12 | 0.47 | 0.6399 | 0.37 | 1.48 | 0.1431 |
| **L CR** | −0.19 | −0.75 | 0.4563 | −0.18 | −0.71 | 0.4813 | 0.15 | 0.61 | 0.5415 | 0.36 | 1.46 | 0.1494 |
| **L CGH** | −0.18 | −0.71 | 0.4778 | −0.18 | −0.71 | 0.4785 | 0.28 | 1.13 | 0.2625 | −0.12 | −0.49 | 0.6271 |
| **R CGC** | −0.16 | −0.66 | 0.5124 | −0.21 | −0.84 | 0.4015 | 0.15 | 0.62 | 0.5360 | 0.24 | 0.97 | 0.3362 |
| **R RLIC** | −0.15 | −0.59 | 0.5593 | −0.09 | −0.35 | 0.7290 | 0.1 | 0.42 | 0.6785 | 0.16 | 0.65 | 0.5172 |
| **L CST** | −0.14 | −0.58 | 0.5613 | −0.51 | −2.04 | 0.0451 | −0.22 | −0.9 | 0.3734 | −0.22 | −0.91 | 0.3680 |
| **L SFO** | −0.13 | −0.54 | 0.5898 | −0.12 | −0.5 | 0.6208 | 0.10 | 0.41 | 0.6828 | −0.14 | −0.55 | 0.5842 |
| **R FXST** | −0.12 | −0.46 | 0.6435 | −0.01 | −0.03 | 0.9793 | 0.10 | 0.42 | 0.6755 | −0.25 | −1.02 | 0.3136 |
| **R PTR** | −0.11 | −0.43 | 0.6659 | −0.06 | −0.23 | 0.8211 | 0.07 | 0.3 | 0.7654 | 0.24 | 0.97 | 0.3380 |
| **L CGC** | −0.10 | −0.39 | 0.6983 | −0.11 | −0.43 | 0.6686 | 0.12 | 0.48 | 0.6297 | 0.44 | 1.76 | 0.0825 |
| **R EC** | −0.10 | −0.41 | 0.6853 | −0.21 | −0.86 | 0.3934 | 0.20 | 0.81 | 0.4232 | 0.08 | 0.32 | 0.7533 |
| **L FXST** | −0.10 | −0.39 | 0.6982 | −0.03 | −0.13 | 0.8999 | 0.08 | 0.34 | 0.7334 | 0.54 | 2.20 | 0.0313 |
| **R IC** | −0.10 | −0.41 | 0.6859 | 0.02 | 0.08 | 0.9335 | 0.03 | 0.14 | 0.8886 | 0.07 | 0.27 | 0.7884 |
| **SCC** | −0.06 | −0.23 | 0.8198 | −0.09 | −0.38 | 0.7050 | −0.09 | −0.37 | 0.7135 | 0.19 | 0.76 | 0.4513 |
| **R CGH** | −0.03 | −0.11 | 0.9092 | −0.07 | −0.27 | 0.7870 | 0.03 | 0.12 | 0.9022 | 0.02 | 0.06 | 0.9497 |
| **L EC** | −0.03 | −0.11 | 0.9138 | −0.10 | −0.41 | 0.6821 | 0.08 | 0.33 | 0.7423 | 0.01 | 0.04 | 0.9681 |
| **R PCR** | 0 | 0 | 0.9977 | 0.04 | 0.16 | 0.8709 | −0.08 | −0.34 | 0.7342 | 0.24 | 0.98 | 0.3290 |
| **L PLIC** | 0.01 | 0.06 | 0.9528 | 0.16 | 0.64 | 0.5237 | −0.04 | −0.17 | 0.8692 | 0.11 | 0.44 | 0.6618 |
| **R UNC** | 0.04 | 0.15 | 0.8818 | 0.08 | 0.31 | 0.7550 | 0 | −0.01 | 0.9942 | 0.15 | 0.62 | 0.5369 |
| **L PCR** | 0.05 | 0.22 | 0.8260 | 0.06 | 0.26 | 0.7965 | −0.07 | −0.3 | 0.7672 | 0.34 | 1.38 | 0.1720 |
| **L IFO** | 0.09 | 0.38 | 0.7086 | 0.06 | 0.25 | 0.7999 | −0.19 | −0.76 | 0.4518 | −0.42 | −1.69 | 0.0948 |
| **R CST** | 0.12 | 0.49 | 0.6243 | 0.01 | 0.04 | 0.9688 | −0.38 | −1.54 | 0.1283 | 0.02 | 0.09 | 0.9257 |
| **L UNC** | 0.12 | 0.49 | 0.6261 | 0.03 | 0.12 | 0.9071 | −0.06 | −0.23 | 0.8218 | −0.07 | −0.29 | 0.7705 |
| **R SCR** | 0.14 | 0.55 | 0.5849 | 0.13 | 0.54 | 0.5928 | −0.13 | −0.53 | 0.5952 | 0.23 | 0.93 | 0.3546 |
| **R SFO** | 0.18 | 0.74 | 0.4589 | 0.13 | 0.52 | 0.6070 | −0.13 | −0.52 | 0.6031 | −0.05 | −0.19 | 0.8524 |
| **R IFO** | 0.20 | 0.82 | 0.4156 | 0.22 | 0.9 | 0.3693 | −0.21 | −0.83 | 0.4098 | −0.04 | −0.18 | 0.8601 |
| **R PLIC** | 0.26 | 1.06 | 0.2939 | 0.37 | 1.5 | 0.1375 | −0.29 | −1.15 | 0.2526 | 0 | −0.01 | 0.9941 |
| **L SCR** | 0.27 | 1.1 | 0.2757 | 0.23 | 0.94 | 0.3519 | −0.25 | −1.01 | 0.3149 | 0.32 | 1.30 | 0.1966 |

Notes: Results from regression *Model 1*: the effect of patient status on ROIs after adjusting for age, sex and average motion. The ROIs are ordered by $FA_t$ effect size. ROIs that pass FDR threshold, $p \leq 0.0109$, are indicated in bold. *Abbreviations:* ACR: anterior corona radiata, $AD_t$: FW adjusted axial diffusivity, ALIC: anterior limb of internal capsule, Average: average of $FA_t$, $AD_t$, $RD_t$, and FW, respectively. BCC: body of corpus callosum, CC: corpus callosum, CGC: cingulum, CGH: cingulum hippocampal portion, CR: corona radiata, CST: corticospinal tract, EC: external capsule, $FA_t$: FW adjusted fractional anisotropy, FW: free-water, FX: fornix, FXST: fornix stria terminalis, GCC: genu of corpus callosum, IC: internal capsule, IFO: inferior fronto occipital fasciculus, L: Left, PCR: posterior corona radiata, PLIC: posterior limb of internal capsule, PTR: posterior thalamic radiation, R: Right, $RD_t$: FW adjusted Radial diffusivity, RLIC: retrolenticular part of IC, ROI: region of interest, SCC: splenium of corpus callosum, SCR: superior corona radiata, SFO: superior fronto-occipital fasciculus, SLF: superior longitudinal fasciculus, SS: sagittal stratum, UNC: uncinate.

*Table S2:* The Cohen's d effect size of patient-control differences for FA, MD, RD and AD (conventional DTI method).

| | FA | | | AD | | | RD | | | MD | | |
|---|---|---|---|---|---|---|---|---|---|---|---|---|
| **ROI** | **d** | **t** | **p** | **d** | **t** | **p** | **d** | **t** | **p** | **d** | **t** | **p** |
| **R ACR** | −0.88 | −3.55 | **0.0007** | −0.15 | −0.62 | 0.5361 | 0.66 | 2.65 | **0.0100** | 0.42 | 1.69 | 0.0962 |
| **L ACR** | −0.62 | −2.48 | 0.0155 | −0.09 | −0.35 | 0.7304 | 0.52 | 2.09 | 0.0400 | 0.36 | 1.46 | 0.1493 |
| **L SLF** | −0.60 | −2.44 | 0.0174 | 0.08 | 0.34 | 0.7382 | 0.52 | 2.11 | 0.0388 | 0.43 | 1.72 | 0.0894 |
| **Average** | −0.56 | −2.27 | 0.0266 | 0.08 | 0.34 | 0.7372 | 0.48 | 1.93 | 0.0584 | 0.38 | 1.52 | 0.1327 |
| **R CR** | −0.49 | −1.97 | 0.0534 | 0.09 | 0.36 | 0.7196 | 0.44 | 1.77 | 0.0811 | 0.34 | 1.37 | 0.1745 |
| **L SS** | −0.48 | −1.93 | 0.0583 | −0.21 | −0.85 | 0.3966 | 0.43 | 1.73 | 0.0889 | 0.26 | 1.04 | 0.3034 |
| **FX** | −0.43 | −1.72 | 0.0895 | 0.24 | 0.96 | 0.3395 | 0.24 | 0.95 | 0.3453 | 0.24 | 0.96 | 0.3411 |
| **R ALIC** | −0.39 | −1.57 | 0.1215 | −0.16 | −0.66 | 0.5126 | 0.13 | 0.52 | 0.6080 | 0.01 | 0.04 | 0.9680 |
| **L ALIC** | −0.37 | −1.50 | 0.1382 | −0.64 | −2.57 | 0.0125 | 0.01 | 0.02 | 0.9834 | −0.34 | −1.36 | 0.1784 |
| **L PTR** | −0.36 | −1.44 | 0.1559 | −0.07 | −0.29 | 0.7709 | 0.33 | 1.34 | 0.1854 | 0.24 | 0.95 | 0.3455 |
| **L CR** | −0.34 | −1.35 | 0.1804 | 0.20 | 0.82 | 0.4146 | 0.39 | 1.56 | 0.1238 | 0.37 | 1.48 | 0.1433 |
| **BCC** | −0.32 | −1.29 | 0.2020 | 0.06 | 0.23 | 0.8205 | 0.33 | 1.33 | 0.1892 | 0.28 | 1.14 | 0.2583 |
| **L RLIC** | −0.32 | −1.30 | 0.1968 | 0.18 | 0.73 | 0.4656 | 0.36 | 1.46 | 0.1478 | 0.38 | 1.54 | 0.1278 |
| **R SLF** | −0.32 | −1.30 | 0.1966 | 0.03 | 0.11 | 0.9152 | 0.33 | 1.34 | 0.1834 | 0.25 | 0.99 | 0.3250 |
| **CC** | −0.30 | −1.19 | 0.2380 | 0.03 | 0.12 | 0.9060 | 0.36 | 1.46 | 0.1483 | 0.28 | 1.13 | 0.2640 |
| **R SS** | −0.30 | −1.20 | 0.2360 | −0.09 | −0.37 | 0.7099 | 0.24 | 0.97 | 0.3363 | 0.14 | 0.57 | 0.5730 |
| **GCC** | −0.26 | −1.06 | 0.2909 | −0.02 | −0.09 | 0.9298 | 0.36 | 1.46 | 0.1482 | 0.25 | 1.01 | 0.3171 |
| **L IC** | −0.26 | −1.04 | 0.3023 | −0.06 | −0.25 | 0.8027 | 0.18 | 0.74 | 0.4633 | 0.10 | 0.42 | 0.6756 |
| **L FXST** | −0.23 | −0.93 | 0.3541 | 0.24 | 0.99 | 0.3267 | 0.44 | 1.79 | 0.0775 | 0.62 | 2.50 | 0.0147 |
| **L PCR** | −0.22 | −0.89 | 0.3781 | 0.27 | 1.09 | 0.2787 | 0.27 | 1.09 | 0.2776 | 0.33 | 1.33 | 0.1869 |
| **R PCR** | −0.22 | −0.87 | 0.3878 | 0.18 | 0.75 | 0.4580 | 0.20 | 0.81 | 0.4225 | 0.23 | 0.92 | 0.3633 |
| **R RLIC** | −0.20 | −0.80 | 0.4279 | 0.08 | 0.31 | 0.7571 | 0.20 | 0.81 | 0.4183 | 0.21 | 0.85 | 0.4002 |
| **R CGC** | −0.17 | −0.67 | 0.5032 | −0.02 | −0.08 | 0.9385 | 0.29 | 1.16 | 0.2503 | 0.24 | 0.96 | 0.3402 |
| **R EC** | −0.15 | −0.61 | 0.5436 | −0.04 | −0.17 | 0.8631 | 0.13 | 0.51 | 0.6135 | 0.07 | 0.30 | 0.7631 |
| **L CGC** | −0.11 | −0.43 | 0.6696 | 0.11 | 0.46 | 0.6458 | 0.38 | 1.52 | 0.1323 | 0.42 | 1.70 | 0.0939 |
| **R IC** | −0.11 | −0.45 | 0.6516 | 0.12 | 0.50 | 0.6193 | 0.07 | 0.29 | 0.7720 | 0.12 | 0.5 | 0.6173 |
| **L SFO** | −0.11 | −0.43 | 0.6703 | −0.11 | −0.43 | 0.6652 | −0.09 | −0.36 | 0.7175 | −0.12 | −0.49 | 0.6248 |
| **L CGH** | −0.10 | −0.42 | 0.6794 | −0.25 | −1.00 | 0.3225 | −0.03 | −0.14 | 0.8915 | −0.16 | −0.64 | 0.5220 |
| **R PTR** | −0.10 | −0.41 | 0.6811 | 0.14 | 0.58 | 0.5610 | 0.25 | 1.01 | 0.3181 | 0.24 | 0.96 | 0.3381 |
| **R UNC** | −0.06 | −0.23 | 0.8202 | 0.15 | 0.62 | 0.5385 | 0.13 | 0.53 | 0.6010 | 0.17 | 0.67 | 0.5041 |
| **SCC** | −0.05 | −0.21 | 0.8315 | 0.02 | 0.10 | 0.9205 | 0.19 | 0.77 | 0.4434 | 0.15 | 0.6 | 0.5525 |
| **L EC** | −0.03 | −0.11 | 0.9142 | −0.03 | −0.11 | 0.9108 | 0.03 | 0.14 | 0.8914 | 0.01 | 0.06 | 0.9522 |
| **R CGH** | 0.03 | 0.11 | 0.9118 | 0.08 | 0.31 | 0.7583 | 0.07 | 0.28 | 0.7795 | 0.11 | 0.45 | 0.6537 |
| **L CST** | 0.03 | 0.13 | 0.8950 | −0.40 | −1.63 | 0.1073 | −0.16 | −0.65 | 0.5210 | −0.43 | −1.72 | 0.0898 |
| **L PLIC** | 0.04 | 0.14 | 0.8860 | 0.25 | 0.99 | 0.3251 | 0.05 | 0.21 | 0.8327 | 0.19 | 0.78 | 0.4402 |
| **R SCR** | 0.05 | 0.19 | 0.8518 | 0.22 | 0.89 | 0.3771 | 0.14 | 0.56 | 0.5775 | 0.23 | 0.94 | 0.3496 |
| **R FXST** | 0.06 | 0.26 | 0.7958 | −0.01 | −0.03 | 0.9758 | −0.10 | −0.39 | 0.7004 | −0.09 | −0.38 | 0.7065 |
| **L UNC** | 0.07 | 0.28 | 0.7803 | −0.04 | −0.16 | 0.8714 | −0.12 | −0.48 | 0.6294 | −0.11 | −0.45 | 0.6522 |
| **L SCR** | 0.09 | 0.37 | 0.7103 | 0.35 | 1.39 | 0.1679 | 0.13 | 0.54 | 0.5932 | 0.31 | 1.27 | 0.2091 |
| **R SFO** | 0.19 | 0.76 | 0.4483 | 0.11 | 0.46 | 0.6490 | −0.14 | −0.55 | 0.5866 | −0.03 | −0.13 | 0.8983 |
| **R IFO** | 0.25 | 1.00 | 0.3195 | 0.17 | 0.69 | 0.4926 | −0.15 | −0.59 | 0.5544 | 0 | −0.02 | 0.9872 |
| **R PLIC** | 0.27 | 1.11 | 0.2730 | 0.41 | 1.64 | 0.1058 | −0.18 | −0.71 | 0.4805 | 0.14 | 0.58 | 0.5658 |
| **R CST** | 0.29 | 1.16 | 0.2500 | 0.14 | 0.56 | 0.5803 | −0.16 | −0.63 | 0.5313 | −0.01 | −0.02 | 0.9831 |
| **L IFO** | 0.34 | 1.37 | 0.1744 | −0.09 | −0.35 | 0.7273 | −0.38 | −1.53 | 0.1299 | −0.44 | −1.78 | 0.0799 |

*Notes:* Results from regression *Model 1*: the effect of patient status on ROIs after adjusting for age, sex and average motion. The ROIs are ordered by FA effect sizes. ROIs that pass FDR threshold, $p \leq 0.0109$, are indicated in bold. *Abbreviations:* ACR: anterior corona radiata, AD: axial diffusivity, ALIC: anterior limb of internal capsule, BCC: body of corpus callosum, CC: corpus callosum, CGC: cingulum, CGH: cingulum hippocampal portion, CR: corona radiata, CST: corticospinal tract, EC: external capsule, FA: fractional anisotropy, FX: fornix, FXST: fornix stria terminalis, GCC: genu of corpus callosum, IC: internal capsule, IFO: inferior fronto occipital fasciculus, MD: mean diffusivity, PCR: posterior corona radiata, PLIC: posterior limb of internal capsule, PTR: posterior thalamic radiation, RD: radial diffusivity, RLIC: retrolenticular part of IC, ROI: region of interest, SCC: splenium of corpus callosum, SCR: superior corona radiata, SFO: superior fronto-occipital fasciculus, SLF: superior longitudinal fasciculus, SS: sagittal stratum, UNC: uncinate.

*Table S3:* The Cohen's d effect size of CPZ on all ROIs in patients.

| | $FA_t$ | | | $AD_t$ | | | $RD_t$ | | | FW | | |
|---|---|---|---|---|---|---|---|---|---|---|---|---|
| **ROI** | **d** | **t** | **p** | **d** | **t** | **p** | **d** | **t** | **p** | **d** | **t** | **p** |
| **R ALIC** | −0.84 | −2.10 | 0.0455 | −0.87 | −2.18 | 0.0392 | 0.70 | 1.75 | 0.0918 | −0.19 | −0.47 | 0.6450 |
| **L SLF** | −0.54 | −1.35 | 0.1882 | −0.38 | −0.94 | 0.3545 | 0.42 | 1.05 | 0.3016 | −0.15 | −0.36 | 0.7195 |
| **R CR** | −0.52 | −1.31 | 0.2026 | −0.43 | −1.07 | 0.2948 | 0.47 | 1.18 | 0.2484 | −0.31 | −0.77 | 0.4513 |
| **R UNC** | −0.52 | −1.30 | 0.2054 | −0.31 | −0.79 | 0.4397 | 0.42 | 1.06 | 0.3001 | −0.20 | −0.50 | 0.6180 |
| **L CR** | −0.49 | −1.23 | 0.2306 | −0.50 | −1.25 | 0.2213 | 0.49 | 1.21 | 0.2365 | −0.22 | −0.54 | 0.5935 |
| **Average** | −0.48 | −1.20 | 0.2421 | −0.60 | −1.49 | 0.1481 | 0.32 | 0.80 | 0.4289 | −0.25 | −0.62 | 0.5394 |
| **L SCR** | −0.48 | −1.19 | 0.2441 | −0.44 | −1.10 | 0.2805 | 0.44 | 1.10 | 0.2830 | −0.38 | −0.95 | 0.3530 |
| **R SCR** | −0.43 | −1.07 | 0.2958 | −0.36 | −0.91 | 0.3713 | 0.37 | 0.92 | 0.3649 | −0.30 | −0.76 | 0.4541 |
| **L PTR** | −0.42 | −1.05 | 0.3041 | −0.27 | −0.68 | 0.5050 | 0.34 | 0.86 | 0.4003 | −0.41 | −1.03 | 0.3147 |
| **R ACR** | −0.35 | −0.89 | 0.3840 | −0.36 | −0.91 | 0.3728 | 0.36 | 0.89 | 0.3821 | −0.16 | −0.40 | 0.6946 |
| **R IC** | −0.35 | −0.88 | 0.3885 | −0.28 | −0.70 | 0.4919 | 0.25 | 0.62 | 0.5408 | −0.27 | −0.68 | 0.5021 |
| **R PCR** | −0.35 | −0.88 | 0.3852 | −0.21 | −0.53 | 0.5974 | 0.33 | 0.82 | 0.4189 | −0.52 | −1.30 | 0.2056 |
| **SCC** | −0.35 | −0.86 | 0.3961 | −0.25 | −0.63 | 0.5376 | 0.24 | 0.60 | 0.5517 | −0.43 | −1.07 | 0.2938 |
| **CC** | −0.32 | −0.81 | 0.4261 | −0.35 | −0.87 | 0.3939 | 0.26 | 0.65 | 0.5234 | −0.34 | −0.84 | 0.4098 |
| **L IFO** | −0.30 | −0.74 | 0.4664 | −0.13 | −0.33 | 0.7460 | 0.24 | 0.59 | 0.5580 | 0.09 | 0.23 | 0.8209 |
| **L SS** | −0.28 | −0.69 | 0.4959 | −0.32 | −0.80 | 0.4295 | 0.27 | 0.68 | 0.5017 | −0.42 | −1.05 | 0.3032 |
| **L ACR** | −0.26 | −0.65 | 0.5231 | −0.33 | −0.82 | 0.4192 | 0.31 | 0.79 | 0.4393 | 0 | 0 | 0.9989 |
| **R PLIC** | −0.25 | −0.64 | 0.5311 | −0.20 | −0.49 | 0.6279 | 0.21 | 0.52 | 0.6077 | 0.02 | 0.04 | 0.9688 |
| **L CST** | −0.24 | −0.60 | 0.5529 | −0.29 | −0.74 | 0.4677 | 0.35 | 0.88 | 0.3849 | −0.11 | −0.28 | 0.7817 |
| **L PCR** | −0.24 | −0.60 | 0.5527 | −0.24 | −0.59 | 0.5598 | 0.22 | 0.55 | 0.5896 | −0.44 | −1.10 | 0.2834 |
| **R SFO** | −0.24 | −0.61 | 0.5506 | −0.31 | −0.77 | 0.4477 | 0.29 | 0.72 | 0.4802 | −0.46 | −1.16 | 0.2564 |
| **GCC** | −0.23 | −0.59 | 0.5628 | −0.14 | −0.35 | 0.7303 | 0.23 | 0.56 | 0.5780 | −0.01 | −0.02 | 0.9859 |
| **L ALIC** | −0.22 | −0.54 | 0.5908 | −0.32 | −0.79 | 0.4354 | 0.20 | 0.49 | 0.6275 | −0.04 | −0.10 | 0.9248 |
| **BCC** | −0.21 | −0.51 | 0.6116 | −0.36 | −0.89 | 0.3831 | 0.15 | 0.38 | 0.7081 | −0.40 | −1.00 | 0.3275 |
| **R IFO** | −0.20 | −0.51 | 0.6150 | −0.18 | −0.45 | 0.6575 | 0.16 | 0.40 | 0.6940 | −0.32 | −0.80 | 0.4335 |
| **L PLIC** | −0.19 | −0.49 | 0.6302 | −0.29 | −0.73 | 0.4704 | 0.12 | 0.30 | 0.7692 | −0.78 | −1.94 | 0.0640 |
| **R SLF** | −0.16 | −0.39 | 0.7016 | 0.03 | 0.09 | 0.9326 | 0 | 0 | 0.9989 | −0.05 | −0.13 | 0.8973 |
| **L UNC** | −0.15 | −0.37 | 0.7120 | −0.08 | −0.20 | 0.8437 | 0.05 | 0.12 | 0.9061 | −0.10 | −0.25 | 0.8047 |
| **L CGC** | −0.14 | −0.34 | 0.7374 | −0.02 | −0.05 | 0.9624 | 0.19 | 0.48 | 0.6383 | −0.42 | −1.04 | 0.3073 |
| **L IC** | −0.14 | −0.36 | 0.7234 | −0.19 | −0.46 | 0.6464 | 0.07 | 0.17 | 0.8668 | −0.48 | −1.21 | 0.2379 |
| **L FXST** | −0.09 | −0.22 | 0.8276 | −0.08 | −0.19 | 0.8525 | 0.05 | 0.14 | 0.8926 | −0.40 | −1.01 | 0.3215 |
| **L EC** | −0.05 | −0.13 | 0.8960 | −0.10 | −0.24 | 0.8140 | 0.09 | 0.22 | 0.8256 | 0.10 | 0.24 | 0.8088 |
| **R EC** | 0 | 0 | 0.9987 | −0.04 | −0.11 | 0.9139 | 0.01 | 0.02 | 0.9824 | −0.48 | −1.20 | 0.2431 |
| **R SS** | 0.01 | 0.03 | 0.9799 | −0.07 | −0.16 | 0.8719 | −0.01 | −0.02 | 0.9847 | −0.44 | −1.10 | 0.2834 |
| **R CGC** | 0.04 | 0.10 | 0.9215 | 0.12 | 0.30 | 0.7702 | 0.02 | 0.06 | 0.9560 | −0.24 | −0.61 | 0.5504 |
| **L RLIC** | 0.08 | 0.21 | 0.8363 | 0.17 | 0.43 | 0.6718 | −0.15 | −0.37 | 0.7133 | −0.32 | −0.81 | 0.4254 |
| **R PTR** | 0.09 | 0.22 | 0.8297 | 0.05 | 0.12 | 0.9056 | −0.09 | −0.23 | 0.8208 | −0.33 | −0.83 | 0.4168 |
| **R CST** | 0.14 | 0.34 | 0.7358 | 0.02 | 0.05 | 0.9621 | −0.02 | −0.05 | 0.9589 | −0.39 | −0.97 | 0.3419 |
| **L SFO** | 0.14 | 0.36 | 0.7240 | 0.08 | 0.19 | 0.8491 | −0.21 | −0.52 | 0.6080 | −0.43 | −1.08 | 0.2920 |
| **R RLIC** | 0.19 | 0.48 | 0.6333 | 0.21 | 0.53 | 0.5974 | −0.21 | −0.53 | 0.6039 | −0.47 | −1.18 | 0.2484 |
| **FX** | 0.20 | 0.51 | 0.6162 | 0.25 | 0.63 | 0.5320 | −0.12 | −0.29 | 0.7718 | −0.10 | −0.24 | 0.8096 |
| **L CGH** | 0.22 | 0.56 | 0.5825 | 0.16 | 0.39 | 0.6972 | −0.28 | −0.69 | 0.4973 | −0.45 | −1.12 | 0.2727 |
| **R FXST** | 0.29 | 0.72 | 0.4777 | 0.26 | 0.64 | 0.5294 | −0.24 | −0.61 | 0.5464 | −0.43 | −1.07 | 0.2934 |
| **R CGH** | 0.43 | 1.08 | 0.2912 | 0.34 | 0.84 | 0.4094 | −0.46 | −1.15 | 0.2622 | −0.31 | −0.77 | 0.4486 |

*Notes:* Results from regression model that investigates the effect of CPZ on ROIs after adjusting for age, sex and average motion. The ROIs are ordered by $FA_t$ effect size. ROIs that passed the FDR threshold, $p \leq 0.0109$, are indicated in bold. *Abbreviations:* ACR: anterior corona radiata, AD: axial diffusivity, ALIC: anterior limb of internal capsule, BCC: body of corpus callosum, CC: corpus callosum, CGC: cingulum, CGH: cingulum hippocampal portion, CR: corona radiata, CST: corticospinal tract, EC: external capsule, FA: fractional anisotropy, FX: fornix, FXST: fornix stria terminalis, GCC: genu of corpus callosum, IC: internal capsule, IFO: inferior fronto occipital fasciculus, MD: mean diffusivity, PCR: posterior corona radiata, PLIC: posterior limb of internal capsule, PTR: posterior thalamic radiation, RD: radial diffusivity, RLIC: retrolenticular part of IC, ROI: region of interest, SCC: splenium of corpus callosum, SCR: superior corona radiata, SFO: superior fronto-occipital fasciculus, SLF: superior longitudinal fasciculus, SS: sagittal stratum, UNC: uncinate.

*Table S4:* The Cohen's d effect size of total SAPS on all ROIs in patients.

| ROI | $FA_t$ | | | $AD_t$ | | | $RD_t$ | | | FW | | |
|---|---|---|---|---|---|---|---|---|---|---|---|---|
| | d | t | p | d | t | p | d | t | p | d | t | p |
| L CGH | −0.83 | −2.06 | 0.0497 | −0.77 | −1.91 | 0.0672 | 0.69 | 1.72 | 0.0981 | 0.73 | 1.84 | 0.0784 |
| R RLIC | −0.83 | −2.08 | 0.0478 | −0.73 | −1.83 | 0.0794 | 0.78 | 1.95 | 0.0623 | 0.72 | 1.79 | 0.0859 |
| L PTR | −0.79 | −1.99 | 0.0580 | −0.77 | −1.93 | 0.0656 | 0.79 | 1.97 | 0.0601 | 0.80 | 2.00 | 0.0566 |
| L IC | −0.65 | −1.63 | 0.1156 | −0.56 | −1.39 | 0.1769 | 0.65 | 1.63 | 0.1156 | 0.42 | 1.05 | 0.3018 |
| R IC | −0.63 | −1.59 | 0.1253 | −0.45 | −1.13 | 0.2710 | 0.69 | 1.74 | 0.0947 | 0.48 | 1.19 | 0.2451 |
| R SS | −0.62 | −1.55 | 0.1326 | −0.51 | −1.28 | 0.2116 | 0.63 | 1.56 | 0.1303 | 1.09 | 2.73 | 0.0113 |
| GCC | −0.58 | −1.46 | 0.1571 | −0.19 | −0.47 | 0.6395 | 0.59 | 1.47 | 0.1552 | 0.09 | 0.24 | 0.8143 |
| R PTR | −0.58 | −1.44 | 0.1616 | −0.50 | −1.25 | 0.2244 | 0.64 | 1.60 | 0.1229 | 1.20 | 2.99 | **0.0061** |
| L ALIC | −0.57 | −1.42 | 0.1691 | −0.73 | −1.81 | 0.0817 | 0.64 | 1.60 | 0.1222 | −0.15 | −0.37 | 0.7140 |
| R SCR | −0.55 | −1.36 | 0.1849 | −0.51 | −1.26 | 0.2180 | 0.50 | 1.26 | 0.2205 | 0.64 | 1.61 | 0.1209 |
| R CGH | −0.53 | −1.32 | 0.1991 | −0.70 | −1.74 | 0.0942 | 0.64 | 1.60 | 0.1217 | 0.08 | 0.20 | 0.8457 |
| R PCR | −0.53 | −1.32 | 0.2002 | −0.35 | −0.88 | 0.3858 | 0.52 | 1.30 | 0.2049 | 0.84 | 2.09 | 0.0467 |
| L ACR | −0.51 | −1.27 | 0.2148 | −0.59 | −1.48 | 0.1525 | 0.55 | 1.38 | 0.1783 | 0.63 | 1.57 | 0.1289 |
| L RLIC | −0.49 | −1.23 | 0.2294 | −0.47 | −1.17 | 0.2544 | 0.48 | 1.20 | 0.2411 | 0.66 | 1.65 | 0.1121 |
| L CR | −0.48 | −1.21 | 0.2375 | −0.45 | −1.14 | 0.2669 | 0.44 | 1.10 | 0.2818 | 0.82 | 2.05 | 0.0511 |
| L PLIC | −0.48 | −1.19 | 0.2452 | −0.26 | −0.66 | 0.5168 | 0.49 | 1.24 | 0.2281 | 0.27 | 0.67 | 0.5102 |
| R CR | −0.40 | −1.01 | 0.3211 | −0.37 | −0.92 | 0.3646 | 0.41 | 1.02 | 0.3155 | 0.65 | 1.62 | 0.1186 |
| CC | −0.39 | −0.96 | 0.3445 | 0 | 0 | 0.9963 | 0.37 | 0.93 | 0.3603 | 0.34 | 0.84 | 0.4068 |
| L PCR | −0.37 | −0.92 | 0.3675 | −0.27 | −0.68 | 0.5021 | 0.29 | 0.74 | 0.4689 | 0.78 | 1.94 | 0.0634 |
| Average | −0.34 | −0.86 | 0.3969 | −0.21 | −0.54 | 0.5967 | 0.53 | 1.33 | 0.1943 | 0.58 | 1.44 | 0.1614 |
| R PLIC | −0.31 | −0.78 | 0.4420 | −0.14 | −0.35 | 0.7326 | 0.35 | 0.88 | 0.3880 | 0.17 | 0.43 | 0.6684 |
| R SFO | −0.23 | −0.59 | 0.5625 | −0.24 | −0.61 | 0.5474 | 0.25 | 0.62 | 0.5407 | 0.67 | 1.68 | 0.1045 |
| L SCR | −0.21 | −0.52 | 0.6056 | −0.18 | −0.45 | 0.6597 | 0.17 | 0.44 | 0.6655 | 1.00 | 2.50 | 0.0194 |
| BCC | −0.20 | −0.51 | 0.6169 | 0.08 | 0.19 | 0.8525 | 0.17 | 0.41 | 0.6821 | 0.43 | 1.06 | 0.2973 |
| SCC | −0.19 | −0.48 | 0.6322 | 0.08 | 0.20 | 0.8407 | 0.24 | 0.60 | 0.5536 | 0.26 | 0.65 | 0.5210 |
| R ALIC | −0.18 | −0.45 | 0.6574 | −0.18 | −0.45 | 0.6594 | 0.42 | 1.05 | 0.3047 | 0.29 | 0.73 | 0.4724 |
| R CGC | −0.18 | −0.44 | 0.6613 | −0.30 | −0.75 | 0.4596 | 0.21 | 0.52 | 0.6081 | 0.46 | 1.14 | 0.2658 |
| L SFO | −0.16 | −0.39 | 0.6977 | −0.04 | −0.11 | 0.9159 | 0.25 | 0.63 | 0.5363 | 0.19 | 0.47 | 0.6427 |
| L EC | −0.13 | −0.32 | 0.7520 | −0.11 | −0.27 | 0.7892 | 0.12 | 0.29 | 0.7721 | 0.40 | 1.01 | 0.3221 |
| L UNC | −0.08 | −0.19 | 0.8517 | −0.06 | −0.16 | 0.8733 | 0.08 | 0.21 | 0.8365 | 0.49 | 1.22 | 0.2354 |
| L SS | −0.07 | −0.18 | 0.8592 | −0.04 | −0.10 | 0.9177 | 0.07 | 0.18 | 0.8608 | 1.16 | 2.89 | **0.0078** |
| R EC | −0.06 | −0.14 | 0.8915 | −0.04 | −0.10 | 0.9201 | 0.02 | 0.06 | 0.9508 | 0.11 | 0.28 | 0.7814 |
| FX | −0.05 | −0.13 | 0.8956 | 0.08 | 0.2 | 0.8417 | 0.82 | 2.06 | 0.0499 | −0.37 | −0.91 | 0.3698 |
| L FXST | −0.04 | −0.11 | 0.9151 | −0.14 | −0.36 | 0.7224 | 0.07 | 0.17 | 0.8665 | 0.36 | 0.90 | 0.3748 |
| R UNC | −0.04 | −0.10 | 0.9187 | 0.02 | 0.04 | 0.9697 | 0.04 | 0.10 | 0.9195 | 0.79 | 1.97 | 0.0597 |
| R FXST | 0 | 0.01 | 0.9957 | 0.05 | 0.13 | 0.8999 | 0.17 | 0.43 | 0.6710 | 0.33 | 0.83 | 0.4164 |
| L CGC | 0.03 | 0.08 | 0.9385 | 0.07 | 0.17 | 0.8629 | −0.05 | −0.13 | 0.9013 | 0.55 | 1.37 | 0.1838 |
| L IFO | 0.09 | 0.23 | 0.8166 | 0.08 | 0.21 | 0.8353 | −0.26 | −0.65 | 0.5205 | −0.23 | −0.57 | 0.5710 |
| R ACR | 0.11 | 0.29 | 0.7778 | 0.06 | 0.15 | 0.8837 | −0.07 | −0.18 | 0.8547 | 0.47 | 1.18 | 0.2507 |
| R SLF | 0.25 | 0.62 | 0.5403 | 0.36 | 0.89 | 0.3820 | −0.28 | −0.70 | 0.4922 | 0.75 | 1.88 | 0.0713 |
| L SLF | 0.43 | 1.07 | 0.2934 | 0.60 | 1.50 | 0.1463 | −0.50 | −1.26 | 0.2198 | 0.90 | 2.24 | 0.0343 |
| R IFO | 0.55 | 1.38 | 0.1805 | 0.61 | 1.53 | 0.1387 | −0.64 | −1.61 | 0.1200 | 0.31 | 0.78 | 0.4422 |
| L CST | 0.83 | 2.07 | 0.0489 | 0.76 | 1.89 | 0.0706 | −0.85 | −2.12 | 0.044 | 0.01 | 0.03 | 0.9749 |
| R CST | 1.05 | 2.62 | 0.0149 | 0.74 | 1.84 | 0.0779 | −0.69 | −1.72 | 0.0982 | −0.19 | −0.48 | 0.6341 |

*Notes:* Results from regression model that investigates the effect of total SAPS on ROIs after adjusting for age, sex and average motion. The ROIs are ordered by ascending effect sizes for $FA_t$. ROIs that pass FDR threshold, $p \leq 0.0109$, are indicated in bold. *Abbreviations:* ACR: anterior corona radiata, AD: axial diffusivity, ALIC: anterior limb of internal capsule, BCC: body of corpus callosum, CC: corpus callosum, CGC: cingulum, CGH: cingulum hippocampal portion, CR: corona radiata, CST: corticospinal tract, EC: external capsule, FA: fractional anisotropy, FX: fornix, FXST: fornix stria terminalis, GCC: genu of corpus callosum, IC: internal capsule, IFO: inferior fronto occipital fasciculus, MD: mean diffusivity, PCR: posterior corona radiata, PLIC: posterior limb of internal capsule, PTR: posterior thalamic radiation, RD: radial diffusivity, RLIC: retrolenticular part of IC, ROI: region of interest, SCC: splenium of corpus callosum, SCR: superior corona radiata, SFO: superior fronto-occipital fasciculus, SLF: superior longitudinal fasciculus, SS: sagittal stratum, UNC: uncinate.

*Table S5:* The Cohen's d effect size of total SANS on all ROIs in patients.

| ROI | $FA_t$ | | | $AD_t$ | | | $RD_t$ | | | FW | | |
|---|---|---|---|---|---|---|---|---|---|---|---|---|
| | d | t | p | d | t | p | d | t | p | d | t | p |
| R ALIC | −1.14 | −2.84 | **0.0088** | −0.97 | −2.42 | 0.0233 | 1.20 | 2.99 | **0.0061** | 0.29 | 0.72 | 0.4757 |
| R IC | −0.99 | −2.48 | 0.0203 | −0.65 | −1.64 | 0.1142 | 0.96 | 2.39 | 0.0245 | 0.57 | 1.43 | 0.1641 |
| L ALIC | −0.94 | −2.35 | 0.0271 | −0.82 | −2.06 | 0.0498 | 0.89 | 2.22 | 0.0360 | −0.07 | −0.18 | 0.8611 |
| GCC | −0.86 | −2.15 | 0.0411 | −0.27 | −0.68 | 0.5025 | 0.97 | 2.41 | 0.0234 | 0.10 | 0.25 | 0.8070 |
| R CGC | −0.68 | −1.71 | 0.0996 | −0.60 | −1.51 | 0.1448 | 0.69 | 1.71 | 0.0990 | 0.11 | 0.28 | 0.7780 |
| R CGH | −0.65 | −1.62 | 0.1189 | −0.66 | −1.65 | 0.1117 | 0.64 | 1.60 | 0.1215 | −0.44 | −1.10 | 0.2810 |
| L IC | −0.65 | −1.63 | 0.1146 | −0.45 | −1.14 | 0.2670 | 0.64 | 1.61 | 0.1202 | 0.55 | 1.38 | 0.1812 |
| L SCR | −0.65 | −1.62 | 0.1183 | −0.61 | −1.52 | 0.1414 | 0.61 | 1.51 | 0.1424 | 0.27 | 0.67 | 0.5070 |
| L CR | −0.54 | −1.34 | 0.1926 | −0.54 | −1.35 | 0.1903 | 0.52 | 1.30 | 0.2067 | 0.33 | 0.82 | 0.4179 |
| R RLIC | −0.53 | −1.34 | 0.1935 | −0.48 | −1.19 | 0.2439 | 0.52 | 1.30 | 0.2062 | 0.74 | 1.86 | 0.0749 |
| L CGH | −0.51 | −1.27 | 0.2154 | −0.48 | −1.19 | 0.2445 | 0.40 | 1.00 | 0.3251 | −0.24 | −0.60 | 0.5512 |
| R ACR | −0.50 | −1.25 | 0.2231 | −0.55 | −1.37 | 0.1817 | 0.55 | 1.38 | 0.1811 | 0.29 | 0.72 | 0.4810 |
| R CR | −0.48 | −1.21 | 0.2386 | −0.49 | −1.21 | 0.2362 | 0.54 | 1.36 | 0.1858 | 0.31 | 0.77 | 0.4500 |
| R PLIC | −0.48 | −1.20 | 0.2426 | −0.23 | −0.58 | 0.5671 | 0.54 | 1.35 | 0.1883 | 0.44 | 1.10 | 0.2818 |
| L PTR | −0.38 | −0.96 | 0.3451 | −0.49 | −1.22 | 0.2342 | 0.42 | 1.05 | 0.3038 | 0.23 | 0.58 | 0.5678 |
| L RLIC | −0.38 | −0.95 | 0.3515 | −0.39 | −0.99 | 0.3340 | 0.41 | 1.03 | 0.3124 | 0.77 | 1.91 | 0.0672 |
| R SCR | −0.34 | −0.86 | 0.4002 | −0.38 | −0.95 | 0.3520 | 0.37 | 0.93 | 0.3634 | 0.31 | 0.79 | 0.4389 |
| L SFO | −0.31 | −0.77 | 0.4487 | −0.35 | −0.87 | 0.3907 | 0.22 | 0.56 | 0.5793 | 0.05 | 0.13 | 0.8959 |
| R UNC | −0.31 | −0.78 | 0.4400 | −0.38 | −0.96 | 0.3487 | 0.35 | 0.87 | 0.3918 | 0.22 | 0.55 | 0.5860 |
| L ACR | −0.30 | −0.76 | 0.4563 | −0.34 | −0.86 | 0.3984 | 0.31 | 0.78 | 0.4453 | 0.28 | 0.69 | 0.4950 |
| L CGC | −0.29 | −0.72 | 0.4758 | −0.20 | −0.49 | 0.6276 | 0.38 | 0.95 | 0.3495 | 0.45 | 1.12 | 0.2737 |
| L PLIC | −0.29 | −0.74 | 0.4683 | −0.07 | −0.17 | 0.8630 | 0.38 | 0.96 | 0.3486 | 0.38 | 0.96 | 0.3472 |
| R SFO | −0.25 | −0.62 | 0.5401 | −0.32 | −0.80 | 0.4288 | 0.30 | 0.76 | 0.4557 | 0.16 | 0.39 | 0.6964 |
| R IFO | −0.22 | −0.55 | 0.5841 | −0.18 | −0.45 | 0.6572 | 0.16 | 0.39 | 0.6996 | −0.20 | −0.49 | 0.6299 |
| L UNC | −0.22 | −0.55 | 0.5854 | −0.29 | −0.73 | 0.4693 | 0.22 | 0.54 | 0.5915 | 0.07 | 0.17 | 0.8639 |
| R CST | −0.16 | −0.39 | 0.6985 | −0.29 | −0.73 | 0.4712 | 0.12 | 0.3 | 0.7646 | −0.79 | −1.97 | 0.0601 |
| L IFO | −0.15 | −0.39 | 0.7017 | −0.13 | −0.32 | 0.7541 | 0.19 | 0.48 | 0.6321 | 0.03 | 0.07 | 0.9444 |
| L CST | −0.14 | −0.35 | 0.7301 | −0.29 | −0.71 | 0.4814 | −0.13 | −0.32 | 0.7522 | −0.80 | −2.00 | 0.0565 |
| R PCR | −0.14 | −0.35 | 0.7295 | −0.12 | −0.30 | 0.7651 | 0.28 | 0.70 | 0.4931 | 0.27 | 0.67 | 0.5116 |
| L FXST | −0.13 | −0.32 | 0.7531 | 0.01 | 0.03 | 0.9769 | 0.10 | 0.25 | 0.8058 | 0.97 | 2.43 | 0.0225 |
| FX | −0.09 | −0.21 | 0.8320 | 0.44 | 1.09 | 0.284 | 0.82 | 2.04 | 0.0516 | −0.32 | −0.80 | 0.4323 |
| CC | −0.08 | −0.20 | 0.8408 | 0.56 | 1.41 | 0.1711 | 0.44 | 1.11 | 0.2770 | −0.23 | −0.58 | 0.5682 |
| SCC | −0.05 | −0.13 | 0.8943 | 0.49 | 1.23 | 0.2293 | 0.47 | 1.17 | 0.2521 | −0.02 | −0.04 | 0.9664 |
| R PTR | −0.04 | −0.11 | 0.9139 | −0.02 | −0.05 | 0.9602 | 0.08 | 0.19 | 0.8511 | 0.15 | 0.37 | 0.7179 |
| L EC | 0.03 | 0.08 | 0.9365 | 0.10 | 0.25 | 0.8072 | −0.07 | −0.18 | 0.8614 | 0.60 | 1.51 | 0.1434 |
| Average | 0.05 | 0.13 | 0.8951 | 0.17 | 0.42 | 0.6815 | 0.11 | 0.28 | 0.7802 | 0.47 | 1.18 | 0.2498 |
| L PCR | 0.05 | 0.12 | 0.9059 | 0.02 | 0.06 | 0.9555 | −0.03 | −0.07 | 0.9416 | 0.43 | 1.09 | 0.2882 |
| R FXST | 0.06 | 0.14 | 0.8876 | 0.27 | 0.67 | 0.5074 | 0.01 | 0.03 | 0.9768 | 0.10 | 0.25 | 0.8051 |
| L SS | 0.11 | 0.26 | 0.7936 | 0.13 | 0.31 | 0.7566 | −0.10 | −0.26 | 0.7964 | 0.28 | 0.69 | 0.4959 |
| BCC | 0.20 | 0.51 | 0.6156 | 0.9 | 2.24 | 0.0344 | 0.04 | 0.10 | 0.9243 | −0.45 | −1.13 | 0.2699 |
| R SS | 0.25 | 0.63 | 0.5343 | 0.31 | 0.77 | 0.4474 | −0.30 | −0.74 | 0.4659 | 0.09 | 0.21 | 0.8319 |
| R EC | 0.38 | 0.94 | 0.3560 | 0.53 | 1.31 | 0.2010 | −0.50 | −1.25 | 0.2219 | 0.34 | 0.85 | 0.4031 |
| L SLF | 0.40 | 1.00 | 0.3254 | 0.51 | 1.27 | 0.2173 | −0.37 | −0.91 | 0.3696 | 0.37 | 0.94 | 0.3584 |
| R SLF | 0.43 | 1.07 | 0.2956 | 0.35 | 0.87 | 0.3902 | −0.44 | −1.09 | 0.2872 | 0.31 | 0.77 | 0.4487 |

*Notes:* Results from regression model that investigates the effect of total SANS on ROIs after adjusting for age, sex and average motion. The ROIs are ordered by ascending effect sizes for $FA_t$. ROIs that pass FDR threshold, $p \leq 0.0109$, are indicated in bold.. *Abbreviations:* ACR: anterior corona radiata, AD: axial diffusivity, ALIC: anterior limb of internal capsule, BCC: body of corpus callosum, CC: corpus callosum, CGC: cingulum, CGH: cingulum hippocampal portion, CR: corona radiata, CST: corticospinal tract, EC: external capsule, FA: fractional anisotropy, FX: fornix, FXST: fornix stria terminalis, GCC: genu of corpus callosum, IC: internal capsule, IFO: inferior fronto occipital fasciculus, MD: mean diffusivity, PCR: posterior corona radiata, PLIC: posterior limb of internal capsule, PTR: posterior thalamic radiation, RD: radial diffusivity, RLIC: retrolenticular part of IC, ROI: region of interest, SCC: splenium of corpus callosum, SCR: superior corona radiata, SFO: superior fronto-occipital fasciculus, SLF: superior longitudinal fasciculus, SS: sagittal stratum, UNC: uncinate.

*Table S6:* Interaction effects between patient status and subcortical structures on the left ACR.

| | **Dependent: Left ACR $FA_t$** | | | | | | **Dependent: Left ACR $AD_t$** | | | | | | **Dependent: Left ACR $RD_t$** | | | | | | **Dependent: Left ACR Free-water** | | | | | |
|---|---|---|---|---|---|---|---|---|---|---|---|---|---|---|---|---|---|---|---|---|---|---|---|---|
| | **Diagnosis** | | | **Diagnosis-by-subcortical structure** | | | **Diagnosis** | | | **Diagnosis-by-subcortical structure** | | | **Diagnosis** | | | **Diagnosis-by-subcortical structure** | | | **Diagnosis** | | | **Diagnosis-by-subcortical structure** | | |
| **Subcortical structure** | **d** | **t** | **p** | **d** | **t** | **p** | **d** | **t** | **p** | **d** | **t** | **p** | **d** | **t** | **p** | **d** | **t** | **p** | **d** | **t** | **p** | **d** | **t** | **p** |
| **L Accumbens** | −0.73 | −2.91 | **0.0049** | 0.25 | 1.02 | 0.3124 | −0.76 | −3.03 | **0.0035** | 0.19 | 0.77 | 0.4453 | 0.73 | 2.88 | **0.0053** | −0.2 | −0.8 | 0.4268 | 0.29 | 1.16 | 0.2517 | −0.19 | −0.78 | 0.4367 |
| **L Amygdala** | −0.72 | −2.83 | **0.0063** | 0.41 | 1.63 | 0.1074 | −0.72 | −2.83 | **0.0062** | 0.4 | 1.59 | 0.1157 | 0.68 | 2.67 | **0.0096** | −0.39 | −1.58 | 0.12 | 0.33 | 1.31 | 0.1955 | −0.48 | −1.92 | 0.0591 |
| **L Caudate** | −0.77 | −3.05 | **0.0034** | 0.29 | 1.15 | 0.2536 | −0.78 | −3.08 | **0.003** | 0.3 | 1.19 | 0.2372 | 0.74 | 2.93 | **0.0046** | −0.3 | −1.22 | 0.226 | 0.43 | 1.7 | 0.0948 | −0.69 | −2.79 | **0.0069** |
| **L Hippocampus** | −0.69 | −2.75 | **0.0077** | 0.77 | 3.11 | **0.0028** | −0.7 | −2.8 | **0.0068** | 0.72 | 2.92 | **0.0048** | 0.66 | 2.63 | **0.0108** | −0.7 | −2.82 | **0.0064** | 0.29 | 1.17 | 0.2456 | −0.36 | −1.44 | 0.1539 |
| **L Pallidum** | −0.74 | −2.93 | **0.0047** | 0.4 | 1.62 | 0.1098 | −0.74 | −2.93 | **0.0046** | 0.38 | 1.53 | 0.1306 | 0.7 | 2.79 | **0.0069** | −0.37 | −1.49 | 0.1414 | 0.4 | 1.59 | 0.1156 | −0.15 | −0.62 | 0.5361 |
| **L Putamen** | −0.84 | −3.31 | **0.0015** | 0.62 | 2.47 | 0.0161 | −0.84 | −3.3 | **0.0016** | 0.6 | 2.41 | 0.0188 | 0.8 | 3.14 | **0.0025** | −0.55 | −2.2 | 0.0312 | 0.49 | 1.94 | 0.0573 | −0.17 | −0.66 | 0.5095 |
| **L Thalamus** | −0.85 | −3.34 | **0.0014** | 0.5 | 2.02 | 0.0481 | −0.86 | −3.39 | **0.0012** | 0.42 | 1.69 | 0.0954 | 0.82 | 3.24 | **0.0019** | −0.43 | −1.7 | 0.0938 | 0.43 | 1.71 | 0.0917 | −0.69 | −2.76 | **0.0075** |
| **L Ventricle** | −0.73 | −2.9 | **0.005** | −0.44 | −1.76 | 0.0831 | −0.75 | −3 | **0.0038** | −0.37 | −1.5 | 0.1388 | 0.72 | 2.86 | **0.0057** | 0.34 | 1.37 | 0.1754 | 0.3 | 1.19 | 0.2374 | −0.43 | −1.75 | 0.085 |
| **R Accumbens** | −0.65 | −2.57 | 0.0124 | 0.11 | 0.44 | 0.6649 | −0.68 | −2.68 | **0.0092** | 0.02 | 0.09 | 0.9251 | 0.63 | 2.5 | 0.015 | −0.06 | −0.24 | 0.8107 | 0.24 | 0.94 | 0.3522 | −0.39 | −1.56 | 0.1233 |
| **R Amygdala** | −0.75 | −2.98 | **0.004** | 0.67 | 2.72 | **0.0084** | −0.76 | −3.01 | **0.0037** | 0.63 | 2.53 | 0.0139 | 0.71 | 2.84 | **0.006** | −0.61 | −2.45 | 0.017 | 0.32 | 1.28 | 0.2062 | −0.41 | −1.65 | 0.1047 |
| **R Caudate** | −0.78 | −3.09 | **0.003** | 0.41 | 1.66 | 0.1026 | −0.79 | −3.13 | **0.0027** | 0.43 | 1.72 | 0.0908 | 0.75 | 2.97 | **0.0042** | −0.43 | −1.7 | 0.0934 | 0.42 | 1.64 | 0.1064 | −0.71 | −2.86 | **0.0057** |
| **R Hippocampus** | −0.66 | −2.64 | **0.0104** | 0.69 | 2.79 | **0.0068** | −0.68 | −2.7 | **0.0088** | 0.61 | 2.46 | 0.0165 | 0.64 | 2.54 | **0.0135** | −0.61 | −2.45 | 0.0172 | 0.28 | 1.1 | 0.2747 | −0.37 | −1.48 | 0.1428 |
| **R Pallidum** | −0.77 | −3.07 | **0.0031** | 0.31 | 1.27 | 0.2095 | −0.77 | −3.07 | **0.0032** | 0.27 | 1.1 | 0.2743 | 0.73 | 2.92 | **0.0048** | −0.26 | −1.03 | 0.3073 | 0.42 | 1.66 | 0.1012 | −0.41 | −1.66 | 0.1023 |
| **R Putamen** | −0.93 | −3.65 | **5e-04** | 0.61 | 2.44 | 0.0175 | −0.92 | −3.63 | **6e-04** | 0.57 | 2.27 | 0.0263 | 0.87 | 3.45 | **0.001** | −0.52 | −2.1 | 0.0398 | 0.53 | 2.08 | 0.0414 | −0.38 | −1.51 | 0.1364 |
| **R Thalamus** | −0.8 | −3.18 | **0.0022** | 0.68 | 2.75 | **0.0076** | −0.82 | −3.25 | **0.0018** | 0.67 | 2.72 | **0.0084** | 0.77 | 3.06 | **0.0032** | −0.66 | −2.66 | **0.0098** | 0.34 | 1.33 | 0.1873 | −0.43 | −1.73 | 0.0891 |
| **R Ventricle** | −0.75 | −2.98 | **0.0041** | −0.41 | −1.64 | 0.1059 | −0.78 | −3.12 | **0.0027** | −0.39 | −1.58 | 0.118 | 0.75 | 2.97 | **0.0042** | 0.39 | 1.58 | 0.1188 | 0.24 | 0.95 | 0.344 | −0.29 | −1.16 | 0.2506 |

*Notes:* Results of *Model* for the effect of interaction between patient and subcortical structures on the left ACR. Structures passing FDR threshold $p \leq 0.0116$are indicated in bold. *Abbreviations*: $AD_t$: FW adjusted axial diffusivity, ACR: anterior corona radiata, $FA_t$: FW adjusted fractional anisotropy, FW: free-water, L: Left, $RD_t$: FW adjusted radial diffusivity, R: Right.

*Table S7:* Interaction effects between patient status and subcortical structures on the right ACR.

| | **Dependent: Right ACR $FA_t$** | | | | | | **Dependent: Right ACR $AD_t$** | | | | | | **Dependent: Right ACR $RD_t$** | | | | | | **Dependent: Right ACR Free-water** | | | | | |
|---|---|---|---|---|---|---|---|---|---|---|---|---|---|---|---|---|---|---|---|---|---|---|---|---|
| | **Diagnosis** | | | **Diagnosis-by-subcortical structure** | | | **Diagnosis** | | | **Diagnosis-by-subcortical structure** | | | **Diagnosis** | | | **Diagnosis-by-subcortical structure** | | | **Diagnosis** | | | **Diagnosis-by-subcortical structure** | | |
| **Subcortical structure** | **d** | **t** | **p** | **d** | **t** | **p** | **d** | **t** | **p** | **d** | **t** | **p** | **d** | **t** | **p** | **d** | **t** | **p** | **d** | **t** | **p** | **d** | **t** | **p** |
| **L Accumbens** | −0.86 | −3.42 | **0.0011** | 0.2 | 0.79 | 0.4305 | −0.84 | −3.35 | **0.0013** | 0.1 | 0.41 | 0.6836 | 0.85 | 3.36 | **0.0013** | −0.1 | −0.41 | 0.6825 | 0.35 | 1.39 | 0.1699 | −0.11 | −0.46 | 0.6497 |
| **L Amygdala** | −1.01 | −3.97 | **2e-04** | 0.19 | 0.76 | 0.4489 | −0.97 | −3.82 | **3e-04** | 0.13 | 0.5 | 0.6169 | 0.97 | 3.83 | **3e-04** | −0.14 | −0.56 | 0.5789 | 0.43 | 1.7 | 0.0943 | −0.54 | −2.16 | 0.0343 |
| **L Caudate** | −1.02 | −4.04 | **1e-04** | 0.45 | 1.81 | 0.0753 | −0.99 | −3.93 | **2e-04** | 0.43 | 1.73 | 0.0891 | 0.99 | 3.94 | **2e-04** | −0.45 | −1.83 | 0.0723 | 0.48 | 1.91 | 0.06 | −0.67 | −2.69 | **0.0091** |
| **L Hippocampus** | −0.92 | −3.64 | **5e-04** | 0.42 | 1.68 | 0.0975 | −0.89 | −3.54 | **8e-04** | 0.32 | 1.28 | 0.2034 | 0.89 | 3.54 | **7e-04** | −0.32 | −1.3 | 0.1969 | 0.39 | 1.55 | 0.1263 | −0.36 | −1.43 | 0.1568 |
| **L Pallidum** | −1 | −3.98 | **2e-04** | 0.4 | 1.62 | 0.1104 | −0.96 | −3.81 | **3e-04** | 0.39 | 1.55 | 0.1252 | 0.96 | 3.82 | **3e-04** | −0.39 | −1.58 | 0.1196 | 0.46 | 1.81 | 0.0743 | −0.23 | −0.91 | 0.3644 |
| **L Putamen** | −1.14 | −4.49 | **0** | 0.25 | 0.99 | 0.3254 | −1.07 | −4.23 | **1e-04** | 0.19 | 0.76 | 0.4514 | 1.08 | 4.26 | **1e-04** | −0.18 | −0.72 | 0.4729 | 0.59 | 2.33 | 0.0229 | −0.12 | −0.47 | 0.6421 |
| **L Thalamus** | −1.06 | −4.18 | **1e-04** | 0.51 | 2.05 | 0.0443 | −1.02 | −4.02 | **2e-04** | 0.47 | 1.88 | 0.0649 | 1.03 | 4.05 | **1e-04** | −0.48 | −1.9 | 0.0619 | 0.54 | 2.14 | 0.036 | −0.56 | −2.25 | 0.0278 |
| **L Ventricle** | −0.95 | −3.79 | **3e-04** | −0.2 | −0.81 | 0.4193 | −0.95 | −3.78 | **3e-04** | −0.17 | −0.68 | 0.4982 | 0.95 | 3.78 | **3e-04** | 0.16 | 0.64 | 0.5273 | 0.36 | 1.43 | 0.1569 | −0.35 | −1.41 | 0.1638 |
| **R Accumbens** | −0.82 | −3.25 | **0.0018** | 0.09 | 0.35 | 0.7296 | −0.8 | −3.17 | **0.0023** | 0 | −0.02 | 0.9845 | 0.8 | 3.17 | **0.0023** | −0.01 | −0.04 | 0.9711 | 0.33 | 1.31 | 0.1947 | −0.47 | −1.88 | 0.0644 |
| **R Amygdala** | −0.96 | −3.81 | **3e-04** | 0.49 | 1.96 | 0.0537 | −0.92 | −3.67 | **5e-04** | 0.41 | 1.67 | 0.1005 | 0.93 | 3.68 | **5e-04** | −0.42 | −1.68 | 0.0983 | 0.4 | 1.6 | 0.1144 | −0.45 | −1.82 | 0.0736 |
| **R Caudate** | −1.01 | −3.99 | **2e-04** | 0.39 | 1.55 | 0.1257 | −0.99 | −3.9 | **2e-04** | 0.37 | 1.5 | 0.1387 | 0.99 | 3.91 | **2e-04** | −0.4 | −1.61 | 0.1113 | 0.48 | 1.91 | 0.0603 | −0.85 | −3.39 | **0.0012** |
| **R Hippocampus** | −0.88 | −3.49 | **9e-04** | 0.53 | 2.14 | 0.0361 | −0.85 | −3.39 | **0.0012** | 0.43 | 1.72 | 0.0893 | 0.85 | 3.39 | **0.0012** | −0.44 | −1.76 | 0.0824 | 0.37 | 1.46 | 0.1482 | −0.44 | −1.76 | 0.0835 |
| **R Pallidum** | −1.04 | −4.14 | **1e-04** | 0.28 | 1.14 | 0.2571 | −1 | −3.96 | **2e-04** | 0.24 | 0.98 | 0.3301 | 1 | 3.97 | **2e-04** | −0.26 | −1.03 | 0.3068 | 0.48 | 1.92 | 0.0587 | −0.5 | −2.02 | 0.047 |
| **R Putamen** | −1.14 | −4.51 | **0** | 0.18 | 0.72 | 0.472 | −1.08 | −4.25 | **1e-04** | 0.12 | 0.47 | 0.6428 | 1.08 | 4.26 | **1e-04** | −0.11 | −0.45 | 0.6541 | 0.62 | 2.46 | 0.0166 | −0.28 | −1.12 | 0.2667 |
| **R Thalamus** | −1 | −3.98 | **2e-04** | 0.51 | 2.04 | 0.0458 | −0.98 | −3.89 | **2e-04** | 0.5 | 2.03 | 0.046 | 0.98 | 3.9 | **2e-04** | −0.51 | −2.07 | 0.0427 | 0.42 | 1.65 | 0.1029 | −0.55 | −2.21 | 0.0309 |
| **R Ventricle** | −0.94 | −3.72 | **4e-04** | −0.03 | −0.14 | 0.8905 | −0.93 | −3.71 | **4e-04** | −0.03 | −0.13 | 0.8958 | 0.93 | 3.71 | **4e-04** | 0.04 | 0.14 | 0.8881 | 0.3 | 1.2 | 0.2361 | −0.17 | −0.69 | 0.4929 |

*Notes:* Results of *Model 2* for the effect of interaction between patient and subcortical structures on the right ACR. Structures passing FDR threshold $p \leq 0.0116$ are indicated in bold. *Abbreviations*: $AD_t$: FW adjusted axial diffusivity, ACR: anterior corona radiata, $FA_t$: FW adjusted fractional anisotropy, FW: free-water, L: Left, $RD_t$: FW adjusted radial diffusivity, R: Right.

*Table S8:* Interaction effects between patient status and subcortical structures on the left ALIC.

| | **Dependent: Left ALIC $FA_t$** | | | | | | **Dependent: Left ALIC $AD_t$** | | | | | | **Dependent: Left ALIC $RD_t$** | | | | | | **Dependent: Left ALIC Free-water** | | | | | |
|---|---|---|---|---|---|---|---|---|---|---|---|---|---|---|---|---|---|---|---|---|---|---|---|---|
| | **Diagnosis** | | | **Diagnosis-by-subcortical structure** | | | **Diagnosis** | | | **Diagnosis-by-subcortical structure** | | | **Diagnosis** | | | **Diagnosis-by-subcortical structure** | | | **Diagnosis** | | | **Diagnosis-by-subcortical structure** | | |
| **Subcortical structure** | **d** | **t** | **p** | **d** | **t** | **p** | **d** | **t** | **p** | **d** | **t** | **p** | **d** | **t** | **p** | **d** | **t** | **p** | **d** | **t** | **p** | **d** | **t** | **p** |
| **L Accumbens** | −0.7 | −2.8 | **0.0068** | −0.14 | −0.56 | 0.5742 | −0.73 | −2.91 | **0.0049** | −0.1 | −0.4 | 0.6877 | 0.71 | 2.83 | **0.0062** | 0.13 | 0.51 | 0.6092 | −0.39 | −1.55 | 0.1262 | 0.07 | 0.28 | 0.7828 |
| **L Amygdala** | −0.66 | −2.61 | **0.0113** | 0.13 | 0.51 | 0.6147 | −0.66 | −2.62 | **0.0109** | 0.19 | 0.75 | 0.4552 | 0.66 | 2.6 | **0.0116** | −0.08 | −0.32 | 0.7529 | −0.31 | −1.21 | 0.2315 | −0.15 | −0.59 | 0.5596 |
| **L Caudate** | −0.77 | −3.04 | **0.0034** | 0.19 | 0.78 | 0.441 | −0.8 | −3.2 | **0.0021** | 0.3 | 1.19 | 0.2373 | 0.77 | 3.06 | **0.0032** | −0.25 | −0.99 | 0.3263 | −0.36 | −1.45 | 0.1527 | −0.56 | −2.28 | 0.0262 |
| **L Hippocampus** | −0.68 | −2.69 | **0.0091** | 0.18 | 0.71 | 0.4778 | −0.7 | −2.77 | **0.0072** | 0.2 | 0.82 | 0.4179 | 0.69 | 2.73 | **0.0082** | −0.18 | −0.72 | 0.4744 | −0.36 | −1.41 | 0.1624 | −0.16 | −0.65 | 0.5162 |
| **L Pallidum** | −0.7 | −2.79 | **0.0069** | 0.09 | 0.38 | 0.7085 | −0.7 | −2.78 | **0.0071** | 0.12 | 0.49 | 0.6232 | 0.69 | 2.75 | **0.0077** | −0.09 | −0.36 | 0.7186 | −0.33 | −1.33 | 0.1882 | −0.19 | −0.76 | 0.4497 |
| **L Putamen** | −0.61 | −2.42 | **0.0183** | 0.36 | 1.46 | 0.1495 | −0.63 | −2.49 | 0.0153 | 0.44 | 1.77 | 0.0808 | 0.62 | 2.45 | 0.0169 | −0.34 | −1.35 | 0.1823 | −0.25 | −0.99 | 0.3276 | 0.07 | 0.29 | 0.7691 |
| **L Thalamus** | −0.67 | −2.65 | **0.0102** | 0.09 | 0.36 | 0.7173 | −0.69 | −2.71 | **0.0087** | 0.12 | 0.48 | 0.6338 | 0.69 | 2.71 | **0.0087** | −0.1 | −0.42 | 0.6765 | −0.29 | −1.15 | 0.2557 | −0.16 | −0.65 | 0.5182 |
| **L Ventricle** | −0.87 | −3.44 | **0.001** | −0.11 | −0.43 | 0.6718 | −0.91 | −3.62 | **6e-04** | −0.06 | −0.24 | 0.8098 | 0.93 | 3.71 | **4e-04** | 0.12 | 0.5 | 0.622 | −0.43 | −1.72 | 0.0896 | −0.28 | −1.14 | 0.2605 |
| **R Accumbens** | −0.69 | −2.74 | **0.008** | 0.12 | 0.48 | 0.6326 | −0.73 | −2.89 | **0.0053** | 0.18 | 0.71 | 0.4822 | 0.68 | 2.69 | **0.009** | −0.09 | −0.37 | 0.7141 | −0.44 | −1.75 | 0.0856 | 0.07 | 0.28 | 0.7838 |
| **R Amygdala** | −0.68 | −2.72 | **0.0083** | 0.11 | 0.45 | 0.6512 | −0.72 | −2.85 | **0.0058** | 0.15 | 0.61 | 0.5454 | 0.69 | 2.75 | **0.0076** | −0.08 | −0.33 | 0.7426 | −0.38 | −1.49 | 0.1406 | −0.02 | −0.08 | 0.9348 |
| **R Caudate** | −0.75 | −2.95 | **0.0045** | 0.07 | 0.28 | 0.7821 | −0.78 | −3.08 | **0.0031** | 0.14 | 0.56 | 0.5757 | 0.76 | 2.99 | **0.0039** | −0.11 | −0.43 | 0.6686 | −0.37 | −1.47 | 0.1456 | −0.65 | −2.62 | **0.0111** |
| **R Hippocampus** | −0.66 | −2.63 | **0.0108** | 0.22 | 0.91 | 0.3678 | −0.7 | −2.77 | **0.0073** | 0.21 | 0.83 | 0.408 | 0.68 | 2.69 | **0.0091** | −0.24 | −0.97 | 0.3352 | −0.36 | −1.44 | 0.1541 | −0.14 | −0.56 | 0.576 |
| **R Pallidum** | −0.71 | −2.84 | **0.0061** | 0.01 | 0.06 | 0.9562 | −0.72 | −2.86 | **0.0056** | 0.06 | 0.23 | 0.8174 | 0.71 | 2.82 | **0.0064** | −0.01 | −0.04 | 0.9697 | −0.32 | −1.27 | 0.2101 | −0.25 | −1.01 | 0.3146 |
| **R Putamen** | −0.68 | −2.67 | **0.0096** | 0.3 | 1.19 | 0.2375 | −0.7 | −2.76 | **0.0075** | 0.39 | 1.56 | 0.1246 | 0.68 | 2.7 | **0.009** | −0.28 | −1.11 | 0.2696 | −0.28 | −1.11 | 0.2705 | −0.16 | −0.64 | 0.5251 |
| **R Thalamus** | −0.68 | −2.71 | **0.0087** | 0.01 | 0.03 | 0.9736 | −0.7 | −2.78 | **0.0072** | 0 | 0 | 0.9985 | 0.69 | 2.75 | **0.0078** | −0.01 | −0.03 | 0.977 | −0.36 | −1.44 | 0.1551 | −0.17 | −0.68 | 0.497 |
| **R Ventricle** | −0.81 | −3.21 | **0.0021** | −0.31 | −1.26 | 0.2119 | −0.84 | −3.33 | **0.0014** | −0.25 | −0.99 | 0.3247 | 0.85 | 3.38 | **0.0012** | 0.35 | 1.41 | 0.1632 | −0.44 | −1.75 | 0.0851 | −0.35 | −1.39 | 0.1687 |

*Notes:* Results of *Model 2* for the effect of interaction between patient and subcortical structures on the left ALIC. Structures passing FDR threshold $p \leq 0.0116$ are indicated in bold. *Abbreviations*: $AD_t$: FW adjusted axial diffusivity, ALIC: anterior limb of internal capsule, $FA_t$: FW adjusted fractional anisotropy, FW: free-water, L: Left, $RD_t$: FW adjusted radial diffusivity, R: Right.

*Table S9:* Interaction effects between patient status and subcortical structures on the fornix.

| | **Dependent: Fornix $FA_t$** | | | | | | **Dependent: Fornix $AD_t$** | | | | | | **Dependent: Fornix $RD_t$** | | | | | | **Dependent: Fornix Free-water** | | | | | |
|---|---|---|---|---|---|---|---|---|---|---|---|---|---|---|---|---|---|---|---|---|---|---|---|---|
| | **Diagnosis** | | | **Diagnosis-by-subcortical structure** | | | **Diagnosis** | | | **Diagnosis-by-subcortical structure** | | | **Diagnosis** | | | **Diagnosis-by-subcortical structure** | | | **Diagnosis** | | | **Diagnosis-by-subcortical structure** | | |
| **Subcortical structure** | **d** | **t** | **p** | **d** | **t** | **p** | **d** | **t** | **p** | **d** | **t** | **p** | **d** | **t** | **p** | **d** | **t** | **p** | **d** | **t** | **p** | **d** | **t** | **p** |
| **L Accumbens** | −0.15 | −0.61 | 0.5419 | 0.19 | 0.78 | 0.4385 | 0.25 | 1.01 | 0.3179 | 0.05 | 0.21 | 0.8337 | 0.76 | 3.04 | **0.0034** | −0.73 | −2.95 | **0.0044** | 0.18 | 0.73 | 0.4651 | 0.04 | 0.16 | 0.877 |
| **L Amygdala** | −0.27 | −1.05 | 0.2961 | 0.05 | 0.22 | 0.8295 | 0.22 | 0.85 | 0.3983 | 0.02 | 0.08 | 0.9335 | 0.86 | 3.41 | **0.0011** | −0.47 | −1.88 | 0.0644 | 0.28 | 1.09 | 0.2808 | 0.11 | 0.44 | 0.6631 |
| **L Caudate** | −0.22 | −0.89 | 0.3765 | −0.07 | −0.3 | 0.7674 | 0.21 | 0.84 | 0.4019 | 0.01 | 0.04 | 0.9672 | 0.81 | 3.23 | **0.0019** | 0 | 0 | 0.9998 | 0.22 | 0.89 | 0.378 | 0.02 | 0.09 | 0.9296 |
| **L Hippocampus** | −0.21 | −0.82 | 0.4124 | 0.08 | 0.33 | 0.7415 | 0.23 | 0.91 | 0.3678 | 0 | 0.02 | 0.9856 | 0.83 | 3.29 | **0.0016** | −0.68 | −2.72 | **0.0083** | 0.21 | 0.85 | 0.3991 | 0.13 | 0.51 | 0.6117 |
| **L Pallidum** | −0.25 | −0.98 | 0.3293 | 0.03 | 0.1 | 0.9175 | 0.19 | 0.76 | 0.452 | 0.01 | 0.05 | 0.9618 | 0.83 | 3.3 | **0.0016** | −0.46 | −1.85 | 0.0689 | 0.28 | 1.09 | 0.2776 | 0.19 | 0.78 | 0.4385 |
| **L Putamen** | −0.34 | −1.35 | 0.1821 | −0.08 | −0.31 | 0.7561 | 0.15 | 0.61 | 0.5465 | 0.02 | 0.08 | 0.9374 | 0.94 | 3.71 | **4e-04** | −0.48 | −1.9 | 0.0615 | 0.34 | 1.33 | 0.1892 | 0.32 | 1.27 | 0.2093 |
| **L Thalamus** | −0.26 | −1.03 | 0.3061 | 0.2 | 0.79 | 0.4321 | 0.22 | 0.87 | 0.3873 | 0.17 | 0.68 | 0.497 | 0.93 | 3.67 | **5e-04** | −0.6 | −2.4 | 0.0192 | 0.25 | 0.99 | 0.3253 | −0.05 | −0.2 | 0.8387 |
| **L Ventricle** | −0.05 | −0.21 | 0.8342 | 0.03 | 0.12 | 0.9061 | 0.44 | 1.75 | 0.0843 | −0.2 | −0.79 | 0.4332 | 0.8 | 3.17 | **0.0023** | 0.32 | 1.3 | 0.197 | 0.03 | 0.14 | 0.8898 | −0.45 | −1.8 | 0.0767 |
| **R Accumbens** | −0.23 | −0.9 | 0.3706 | 0.08 | 0.31 | 0.7563 | 0.2 | 0.81 | 0.4232 | 0.02 | 0.09 | 0.926 | 0.72 | 2.83 | **0.0062** | −0.43 | −1.71 | 0.0913 | 0.29 | 1.15 | 0.2549 | 0.01 | 0.05 | 0.9587 |
| **R Amygdala** | −0.22 | −0.87 | 0.3862 | 0.15 | 0.59 | 0.5557 | 0.24 | 0.95 | 0.347 | 0.11 | 0.45 | 0.6527 | 0.88 | 3.5 | **8e-04** | −0.54 | −2.18 | 0.0331 | 0.21 | 0.83 | 0.4076 | −0.09 | −0.35 | 0.7277 |
| **R Caudate** | −0.23 | −0.9 | 0.3712 | 0.06 | 0.23 | 0.8189 | 0.21 | 0.83 | 0.412 | 0.17 | 0.68 | 0.4993 | 0.82 | 3.25 | **0.0019** | −0.11 | −0.43 | 0.6665 | 0.23 | 0.89 | 0.3763 | −0.07 | −0.28 | 0.7784 |
| **R Hippocampus** | −0.16 | −0.63 | 0.5291 | 0.19 | 0.78 | 0.4393 | 0.28 | 1.09 | 0.2777 | 0.13 | 0.52 | 0.6053 | 0.83 | 3.28 | **0.0017** | −0.66 | −2.65 | **0.01** | 0.16 | 0.64 | 0.5239 | −0.04 | −0.15 | 0.8797 |
| **R Pallidum** | −0.22 | −0.89 | 0.3771 | 0.08 | 0.34 | 0.7357 | 0.22 | 0.87 | 0.3864 | 0.02 | 0.09 | 0.9307 | 0.86 | 3.43 | **0.0011** | −0.5 | −2.02 | 0.048 | 0.23 | 0.92 | 0.3601 | 0.15 | 0.62 | 0.5369 |
| **R Putamen** | −0.36 | −1.4 | 0.1655 | 0.02 | 0.07 | 0.9423 | 0.13 | 0.51 | 0.6129 | 0.13 | 0.53 | 0.5947 | 0.94 | 3.73 | **4e-04** | −0.33 | −1.33 | 0.1889 | 0.33 | 1.28 | 0.2041 | 0.1 | 0.39 | 0.6967 |
| **R Thalamus** | −0.26 | −1.04 | 0.3019 | 0.33 | 1.33 | 0.1872 | 0.2 | 0.81 | 0.4225 | 0.35 | 1.41 | 0.1621 | 0.83 | 3.3 | **0.0016** | −0.32 | −1.29 | 0.1999 | 0.25 | 0.99 | 0.3237 | −0.14 | −0.57 | 0.5678 |
| **R Ventricle** | 0 | 0.01 | 0.9952 | 0.42 | 1.7 | 0.0948 | 0.5 | 2 | 0.05 | −0.01 | −0.03 | 0.9759 | 0.82 | 3.24 | **0.0019** | 0.03 | 0.11 | 0.9143 | −0.06 | −0.26 | 0.7974 | −0.69 | −2.78 | **0.0071** |

*Notes:* Results of *Model 2* for the effect of interaction between patient and subcortical structures on the fornix. Structures passing FDR threshold $p \leq 0.0116$ are indicated in bold.
*Abbreviations*: $AD_t$: FW adjusted axial diffusivity, $FA_t$: FW adjusted fractional anisotropy, FW: free-water, L: Left, $RD_t$: FW adjusted radial diffusivity, R: Right.
